# Chemical rescue of mutant proteins in living cells by naturally occurring small molecules

**DOI:** 10.1101/2021.02.21.432174

**Authors:** Daniel S. Hassell, Marc G. Steingesser, Ashley S. Denney, Courtney R. Johnson, Michael A. McMurray

## Abstract

Intracellular proteins function in a complex milieu wherein small molecules influence protein folding and act as essential cofactors for enzymatic reactions. Thus protein function depends not only on amino acid sequence but also on the concentrations of such molecules, which are subject to wide variation between organisms, metabolic states, and environmental conditions. We previously found evidence that exogenous guanidine reverses the phenotypes of specific budding yeast septin mutants by binding to a WT septin at the former site of an Arg side chain that was lost during fungal evolution. Here we used a combination of targeted and unbiased approaches to look for other cases of “chemical rescue” by naturally occurring small molecules. We report *in vivo* rescue of hundreds of yeast mutants representing a variety of genes, including likely examples of Arg or Lys side chain replacement by the guanidinium ion. Failed rescue of targeted mutants highlight features required for rescue, as well as key differences between the *in vitro* and *in vivo* environments. Some non-Arg mutants rescued by guanidine likely result from “off-target” effects on specific cellular processes in WT cells. Molecules isosteric to guanidine and known to influence protein folding had a range of effects, from essentially none for urea, to rescue of a few mutants by DMSO. Strikingly, the osmolyte trimethylamine-*N*-oxide rescued ∼20% of the mutants we tested, likely reflecting combinations of direct and indirect effects on mutant protein function. Our findings illustrate the potential of natural small molecules as therapeutic interventions and drivers of evolution.

## INTRODUCTION

Physicochemical variability in the environment drives selection for traits that allow organisms to adapt to prevailing conditions with minimal energetic cost. Temperature, hydrostatic pressure, and the concentrations of intracellular molecules strongly influence protein structure and, consequently, function. Whereas protein sequences in extremophiles provide clear signatures of adaptation to temperature, pressure, intracellular pH and salinity (1), many proteins also depend on the presence of other small molecule ligands as folding or enzymatic cofactors that confer changes in tertiary structure or catalytic activity, respectively. Thus the amino acid sequence of a protein is not sufficient to confer proper folding and function in all environments; likewise, which changes in protein sequence affect function depends on the physicochemical environment.

Researchers commonly exploit “conditional” mutant alleles as powerful tools to manipulate gene function in living cells. Temperature-sensitive (TS) mutants typically harbor one or more missense mutations that shift the folding equilibrium such that the mutant protein functions at least somewhat normally at the “permissive” temperature but loses function at the “nonpermissive”/“restrictive” temperature. Other examples of conditional alleles include analog-sensitive kinases, in which substitutions of one or more key residues in the active site of a protein kinase allow only the mutant form to accept an ATP analog that inhibits kinase activity (2). Here the side chains of the replacement residues are carefully chosen to make critical molecular contacts with the inhibitor molecule.

A conceptually similar approach, but with the goal of investigating the molecular details of enzyme function *in vitro*, involves substituting key amino acids for which the side chains are suspected to make critical contacts in the active site, and attempting to restore function by adding small molecules that bind at the molecular vacancy created by the mutation and restore the missing contacts. There are numerous instances in which this approach, often called “chemical rescue” or “chemical complementation”, has been successful (reviewed in (3–5)). However, we are aware of only a handful of cases in which chemical rescue of a mutant protein has been demonstrated *in vivo* using simple, common small molecules (as opposed to complex molecules tailored to a specific mutant protein). The first example of *in vivo* chemical rescue involved a mutant of the protein kinase Csk in which a key Arg residue was mutated to Ala (6). This study followed earlier *in vitro* studies of the same mutant in which the authors found that the activity of the mutant enzyme could be partially restored by the addition of guanidine hydrochloride (GdnHCl) or, with even greater effect, imidazole (7). GdnHCl dissociates to produce the guanidinium ion (Gdm), which mimics the guanidino group at the end of an Arg side chain and had been used successfully in many *in vitro* chemical rescue experiments of Arg-mutant enzymes. It was a surprise that imidazole, which resembles a His side chain and had been used previously to rescue His-mutant enzymes, also rescued Csk(R318A). Within 15 minutes after exposure of mouse embryonic fibroblasts expressing the Csk(R318A) enzyme to 50 mM imidazole, cellular indications of restored Csk activity were observed, though the extent of rescue was estimated as about 5% of WT (7). Recognizing that the Arg in question is highly conserved among protein kinases in the same family as Csk, the same group went on to mutate the equivalent Arg in the protein kinase Src and show that imidazole could also rescue Src(R388A), this time to nearly 50% of WT activity using only 2 mM imidazole (8).

While free imidazole is not commonly found in nature, the choice of this molecule as a rescue agent has been touted as advantageous relative to guanidine, which is often considered to be toxic (3). Likely for this reason, GdnHCl had not (to our knowledge) been tested for its effectiveness in chemical rescue of Arg-mutant proteins *in vivo*. However, there is abundant evidence that GdnHCl can specifically alter protein function without grossly disrupting cellular function. For decades, researchers have added 3-5 mM GdnHCl to the growth medium of the budding yeast *Saccharomyces cerevisiae* to inhibit the function of the disaggregase protein Hsp104, a non-essential protein required for the propagation of most prions (9–11). Gdm binds in the ATP-binding pocket of this AAA+-family ATPase (12). Notably, ≥5 mM GdnHCl cures yeast cells of the prion [*ISP*^+^] in an Hsp104-independent manner, and inhibits growth specifically of [*ISP*^+^] cells (13), providing evidence of additional, Hsp104-independent effects and reinforcing the idea that GdnHCl is well tolerated by WT cells. Studies in cultured mammalian cells showed that exogenous GdnHCl blocks replication of enteroviruses (14, 15), with no adverse cellular effects up to 10 mM (14). Subsequent work pointed to the viral 2C protein as the direct target (16). [That 2C and Hsp104 are both AAA+ ATPases may be a coincidence.] Gdm also binds in the pores of voltage-gated potassium channels and promotes acetylcholine release (17), representing the likely mechanism of action in the treatment of Lambert-Eaton Myasthenic Syndrome and botulism, for which GdnHCl is approved by the Food and Drug Administration of the United States of America. When combined with another drug with a distinct target, low-dose GdnHCl is effective as a treatment of Lambert-Eaton Myasthenic Syndrome, with few side effects (18). Thus, far from being non-specifically toxic, at low concentrations GdnHCl is innocuous or even beneficial, curing yeast cells of some prions and mammalian cells of some viruses, and treating some human disorders.

We were inspired to investigate the potential of GdnHCl for chemical rescue in living cells by our prior, serendipitous discovery that GdnHCl reverses the TS phenotypes of *S. cerevisiae* cells carrying mutations in the *CDC10* gene (19). *CDC10* encodes a subunit of the essential yeast septin protein complex. Our data indicate that, rather than binding to mutant Cdc10 molecules, Gdm binds to WT molecules of another septin protein in the same complex, at the former site of an Arg side chain that was “lost” during fungal evolution. In many yeast species, loss of the Arg and of GTPase activity by the same septin constrained the kinds of septin hetero-oligomers that are able to assemble in those cells (19). Specifically, we think that Gdm alters the conformation of the homodimerization interface that encompasses the GTP-binding pocket in a way that promotes bypass of Cdc10 subunits during septin complex assembly (19). Restoring the “missing” Arg residue was insufficient to recapitulate the effect of GdnHCl, but it blocked the ability of GdnHCl to rescue, consistent with a model in which multiple amino acid changes during evolution disfavored Cdc10 bypass, and Gdm acts as a “chemical chaperone” to influence folding of a single septin protein in ways that cannot be predicted by protein sequence alone. Nonetheless, our findings lent further support to the idea that GdnHCl might be able to restore function to an Arg-mutant protein in living cells.

Guanidine is not unique among naturally occurring small molecules in its ability to rescue mutant enzyme activity *in vitro* by replacing a functional group from a “missing” side chain. For example, the crystal structure of a mutant form of the protease trypsin, in which an active site Asp residue was replaced by Ser, revealed an unexpected acetate ion from the crystallization solution bound in the void created by the mutation (20). Subsequent tests revealed that exogenous acetate partially restored *in vitro* activity of the mutant enzyme (20). In such cases where the exogenous ligand plays a direct role in catalysis, the high entropic cost of properly ordering the ligand in the mutant active site likely constrains the extent of chemical rescue to a fraction of the activity of a WT enzyme, where the functional group(s) are tethered to the polypeptide backbone in the context of a side chain. With this in mind, it is worth noting previous studies of “chemical repair” of Arg-to-Cys-mutant proteins, in which a synthetic compound with a sulfhydryl-reactive group at one end and a guanidino group at the other is attached covalently to the Cys side chain (21). Here some rescue was observed *in vitro*, but these studies could not exclude the possibility that rescue resulted from more general effects of the small molecule acting at other sites on the mutant protein.

Indeed, guanidine itself (22) and other small molecules isosteric with guanidine, including urea (22) and DMSO (23), are also able to interact with proteins rather non-specifically to alter their conformations, driving complete unfolding at sufficiently high concentrations. The effects of urea are particularly relevant *in vivo*. Some marine animals accumulate intracellular urea at high concentrations (∼0.4 M), the denaturing effects of which they counteract by also accumulating methylamine compounds, including trimethylamine *N*-oxide (TMAO) (24). Deep-sea creatures also accumulate TMAO to counteract protein-unfolding effects of hydrostatic pressure (25). TMAO, like DMSO, is also used by some organisms as an alternative electron receptor for respiration in anaerobic conditions (26). Exogenously added TMAO has further been found to restore function to three otherwise misfolded mutant proteins in living *S. cerevisiae* and *E. coli* cells (27, 28) and one in cultured mammalian cells (29). Thus intracellular TMAO and other naturally occurring compounds with similar effects on protein folding may allow the organisms that produce them to explore a larger protein sequence space (30, 31). The recent discoveries that multiple bacterial species evolved ways to sense and export or detoxify intracellular Gdm (32) raise the possibility that Gdm could also influence protein function and evolution in natural settings. However, we are unaware of any thorough comparison of the effects of naturally occurring small molecules on mutant protein function in living cells.

In this study, we tested the ability of GdnHCl to provide chemical rescue to a variety of Arg-mutant proteins in living cells. We also performed an unbiased search for cases of chemical rescue by GdnHCl using collections of TS mutants and then, based on our findings, examined the potential for rescue by urea, DMSO and TMAO. Our findings document what are to our knowledge the first examples of *bona fide* GdnHCl chemical rescue *in vivo*, and provide a number of molecular insights into the “on-target” and “off-target” effects of GdnHCl, as well as mechanistic comparisons with other naturally-occurring small molecules on mutant protein function.

## RESULTS AND DISCUSSION

### *E. coli* β-gal R599A, a mutant enzyme with reduced activity for specific substrates *in vitro*, functions normally *in vivo*

We first tested *in vivo* chemical rescue by GdnHCl using the most obvious candidate from the *in vitro* chemical rescue literature. *E. coli* β-gal is widely used as a reporter of transcription due to its chromogenic activity on specific analogs of lactose, its native substrate. Arg at position 599 makes intramolecular contacts that position the active site in an “open” conformation with high affinity for substrates (33). *In vitro*, binding of β-gal(R599A) to *ortho*- or *para-*nitrophenyl-β-D-galactoside (*o*NPG or *p*NPG) is reduced ∼10-fold relative to WT, and the kinetics of *o*NPG catalysis are reduced ∼5-fold but can be improved ∼2-fold (to ∼2.5-fold less than WT) by the addition of 100-150 mM GdnHCl (33). As a first step towards asking if GdnHCl can rescue β-gal(R599A) activity *in vivo*, we grew *E. coli* cells mutated for the endogenous *lacZ* gene (encoding β-gal) and expressing WT β-gal or β-gal(R599A) from plasmids on solid medium containing β-gal (5-bromo-4-chloro-3-indoyl-β-d-galactopyranoside; neither *o*NPG nor *p*NPG are not suitable for *in vivo* analysis). The R599A mutation had no obvious effect on blue colony color resulting from β-gal cleavage (Fig. S1). To more sensitively monitor enzyme activity *in vivo*, we exposed growing cultures to X-gal and chloramphenicol, to prevent new β-gal synthesis. X-gal cleavage was assessed in aliquots of cells taken at various timepoints thereafter by adding strong base to lyse the cells and denature the enzyme, and quantifying the blue product using a 96-well plate reader. The R599A mutation had no discernable effect on the kinetics of X-gal cleavage *in vivo* (Fig. S1). We confirmed the presence of the R599A mutation by restriction enzyme digestion of plasmid DNA isolated from the same cells at the end of the experiment (Fig. S1).

The strain we used for these experiments, DH5*α*, harbors at the genomic *lacZ* an N-terminally truncated allele that can be functionally complemented by any other allele that contains the missing residues (34). *lac* operon transcription is repressed unless a galactoside is present in the medium (35). While we added no galactoside in our experiments, to rule out the possibility that the truncated *lacZ* allele contributed to the β-gal activity we observed, we introduced the plasmids into a strain carrying a complete deletion of *lacZ*. X-gal cleavage by β-gal(R599A) was indistinguishable from WT (Fig. S1), confirming our findings in DH5*α*.

Our findings with X-gal as an *in vivo* β-gal(R599A) substrate fit with other results from the literature. Arg at the position corresponding to Arg 599 in *E. coli* is not conserved in β-galactosidases from several other closely-related bacterial species, including *Lactobacillus bulgaricus* (Fig. S1), the β-gal-encoding gene of which was originally cloned via *in vivo* X-gal activity upon heterologous expression in *E. coli* (36). *E. coli* ebgA, a paralog of β-gal active towards X-gal *in vivo* (37), also lacks Arg in this position (Fig. S1). Thus, despite the apparent importance of this residue for β-gal activity on *o*NPG *in vitro*, it is not very important for lactose cleavage *in vivo*. Even *in vitro* the effects of the R599A depend on the substrate: in contrast to *o*NPG, the kinetics of *p*NPG catalysis by β-gal(R599A) are actually slightly faster relative to WT (33). These experiments taught us an important lesson: in the search for examples of chemical rescue *in vivo*, focus on mutants for which there is already some evidence of an *in vivo* defect.

### *In vivo* chemical rescue of Arg-mutant ornithine transcarbamylase by GdnHCl

The drastic reduction of *E. coli* ornithine transcarbamylase (EcOTC) activity *in vitro* observed upon substitution to Gly of Arg57, a residue in the active site, can be partially reversed (back to 10% of WT) by the addition of GdnHCl (38). Human OTC (HsOTC) is 38% identical to EcOTC, solved structures of the two proteins are superimposable, and the Arg57 equivalent is found at position 92 (Fig. 1*A,B*). Mutations in HsOTC cause ornithine transcarbamylase deficiency (OTCD), the most frequent inherited defect in the urea cycle (∼1 in 14,000 live births) characterized by hyperammonemia and a requirement for dietary Arg (39). Among known HsOTC mutations linked to OTCD, Arg92 has been found in a number of cases substituted to other amino acids, including Gly (39), indicating that HsOTC(R92G) is likely a loss-of-function allele *in vivo*. We asked if GdnHCl could rescue mutant OTC function in living yeast cells.

**Figure 1.**
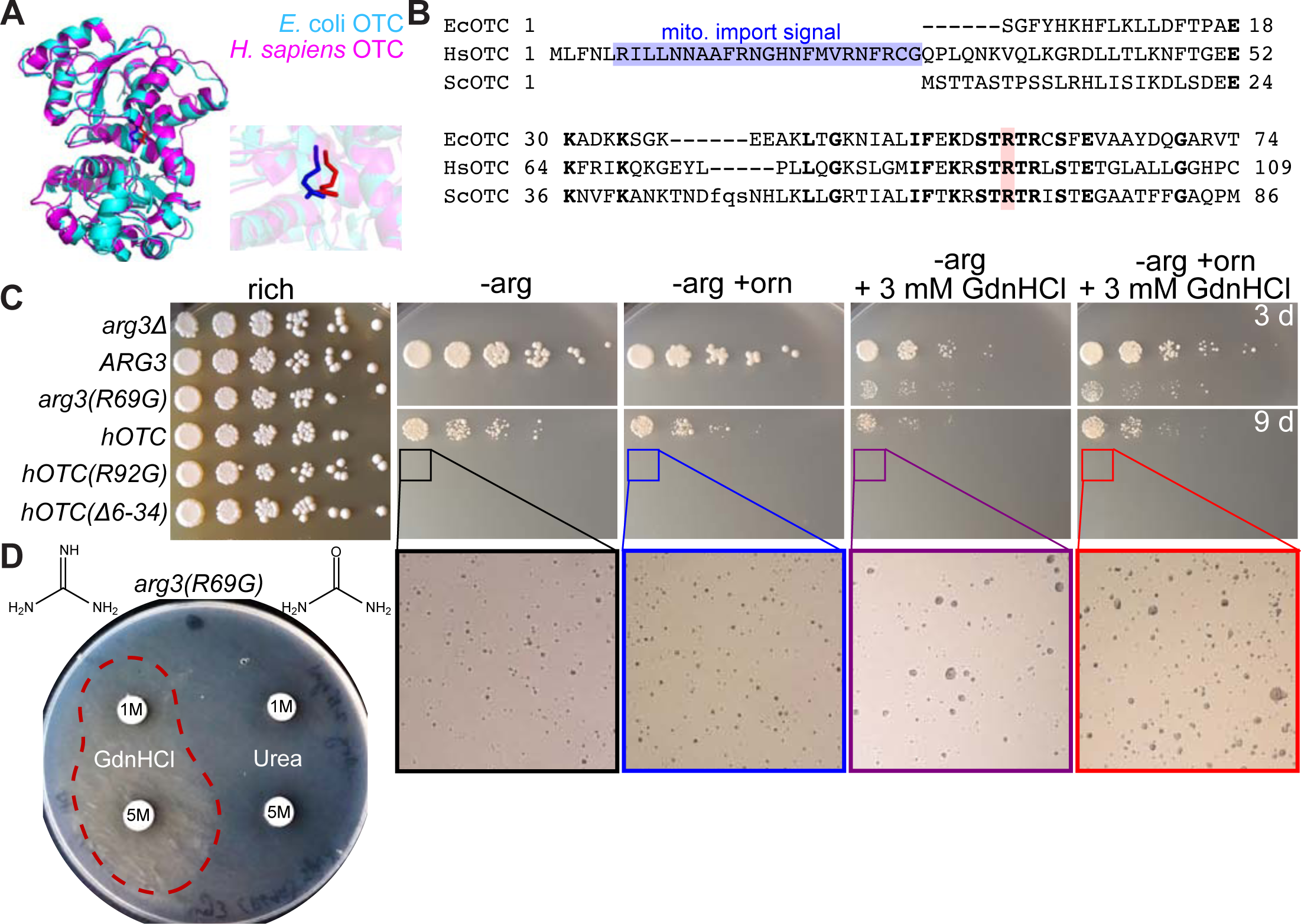
Chemical rescue by GdnHCl of Arg-mutant ornithine transcarbamylase in living yeast cells. (A) Superimposition of crystal structures of human and *E. coli* ornithine transcarbamylase (OTC). The zoomed-in image shows the locations of the side chains of Arg57 from the bacterial enzyme (blue) and Arg92 from the human enzyme. (B) Sequence alignment of portions of the OTC homologs from *E. coli* (EcOTC), human (HsOTC), and *S. cerevisiae* (ScOTC). Bold residues are identical. Blue highlights the leader sequence that directs import of HsOTC into mitochondria. Pink highlights the position of Arg57 in EcOTC, Arg92 in HsOTC, and Arg69 in ScOTC. (C) Dilution series of cells of the indicated genotypes on rich (YPD) or synthetic medium lacking arginine (“-arg”) and thus selective for OTC function. Where indicated, ornithine (“+orn”) and/or GdnHCl were added to final concentration of 100 µg/mL or 3 mM, respectively. Selective plates were incubated at 30°C for the indicated number of days (“3 d” or “9 d”) prior to imaging; YPD plate was incubated for 3 days. (D) A lawn of cells of the indicated genotype was plated to media selective for OTC function and a sterile paper disc soaked with GdnHCl or urea at the indicated concentrations was placed on the lawn. The plate was then imaged after 6 days incubation at 30°C. Strains were: “ARG3”,; “WT recoded”, H06700; “*ura3(R253A)*”, H06701; “*ura3*Δ”, BY4741.

In *S. cerevisiae*, OTC is encoded by the *ARG3* gene, so named because loss-of-function mutants cannot synthesize Arg (40). The Arg3 protein (which we also refer to as “ScOTC”) is 33% identical to EcOTC and is able to complement the Arg auxotrophy of an *E. coli argF* mutant (40). Arg69 in ScOTC corresponds to Arg57 in EcOTC (Fig. 1*B*). We predicted that *arg3(R69G)* mutant yeast would be auxotrophic for Arg and that this auxotrophy would be rescued by the addition of 3 mM GdnHCl to the medium, a concentration that rescues *cdc10* mutants (19). We engineered the R69G substitution in a plasmid-encoded copy of *ARG3* and introduced the mutant plasmid or its WT parent into *arg3*Δ cells. As predicted, Arg prototrophy was conferred by the WT plasmid but not by the R69G mutant unless GdnHCl was present (Fig. 1*C*). Rescue of ScOTC(R69G) function was also observed by placing a disc of GdnHCl-soaked paper onto a lawn of cells (Fig. 1*D*). Notably, an equivalent experiment with urea, which provided weak *in vitro* rescue of EcOTC(R57G) (38), revealed no such effect *in vivo* (Fig. 1*D*). We interpret these data as evidence of partial chemical rescue by GdnHCl of ScOTC(R69G) in living cells.

We wanted to test if GdnHCl could rescue the function of HsOTC(R92G) upon heterologous expression in yeast. WT HsOTC expressed in *arg3* mutant yeast partially complements the Arg auxotrophy (41) (Fig. 1*C*). Adding ornithine to the medium to increase substrate concentration did not increase the extent of complementation (Fig. 1*C*). By contrast, HsOTC(R92G) completely failed to complement (Fig. 1*C*). The addition of 3 mM GdnHCl barely rescued HsOTC(R92G), visualized as slightly larger microcolonies after extended incubation (Fig. 1*C*). Considering the partial GdnHCl rescue of ScOTC(R69G) and the partial complementation by WT HsOTC, our observation of minimal GdnHCl rescue of HsOTC(R92G) was not surprising. We wondered if we could improve the function of HsOTC in yeast, and thereby improve our chances of detecting rescue of HsOTC(R92G). One way HsOTC differs from ScOTC and EcOTC is that it harbors a 32-residue N-terminal leader peptide directing import into mitochondria (Fig. 1*B*). In yeast cells, import of WT HsOTC into the mitochondrial matrix and proteolytic removal of the leader peptide is inefficient, and some HsOTC remains stuck in the mitochondrial membrane (41). Considering that ScOTC is a cytosolic enzyme, citrulline production by HsOTC in the yeast mitochondrion is likely not optimal for *arg3* cells, and HsOTC stuck in the membrane is non-functional (41). We thought that cytosolic HsOTC might generate more usable citrulline. To retain HsOTC in the cytosol, we removed the import sequence, but rather than improving HsOTC function, this alteration eliminated it (Fig. 1*C*). Whereas HsOTC requires (re-)folding assistance from the mitochondrial chaperonin (42), ScOTC requires cytosolic Hsp70 chaperones (43). Thus the poor function of WT HsOTC in yeast and the correspondingly slight rescue by GdnHCl of HsOTC(R92G) likely represent multiple compartment-specific incompatibilities arising from heterologous expression.

### Weak GdnHCl rescue of Arg-mutant orotidine-5’-phosphate decarboxylase, the *URA3* gene product

Due to an exceptionally high affinity for a transition state of its substrate, orotidine-5’-phosphate decarboxylase (ODCase) displays one of the largest rate enhancements (>10^17^-fold) of any known enzyme (44). The guanidino group in the side chain of Arg 235 of the *S. cerevisiae* ODCase, Ura3, contacts the substrate in its transition state (45). *In vitro*, Ura3(R235A) decarboxylates orotidine-5’-phosphate ∼15-fold more slowly than does WT Ura3, but the addition of GdnHCl at 100 mM restores activity to ∼70% of WT (46). To look for rescue by GdnHCl *in vivo*, we introduced the R235A substitution into the *URA3* locus of otherwise WT cells. A guide RNA targeted Cas9 cleavage in the protein-coding sequence, which was repaired by homologous recombination with a synthetic double-stranded DNA molecule encoding WT Ura3 or Ura3(R235A) harboring synonymous substitutions within the sequences between the site of cleavage and codon 235 (Fig. S2). Consistent with the decreased ODCase activity of Ura3(R235A) *in vitro* (46), *ura3(R235A)* cells formed very small colonies on medium lacking uracil (Fig. 2*A*). These cells proliferated slightly faster when 3 mM GdnHCl was added to the media (Fig. 2*A*), indicating weak rescue of ODCase activity.

**Figure 2.**
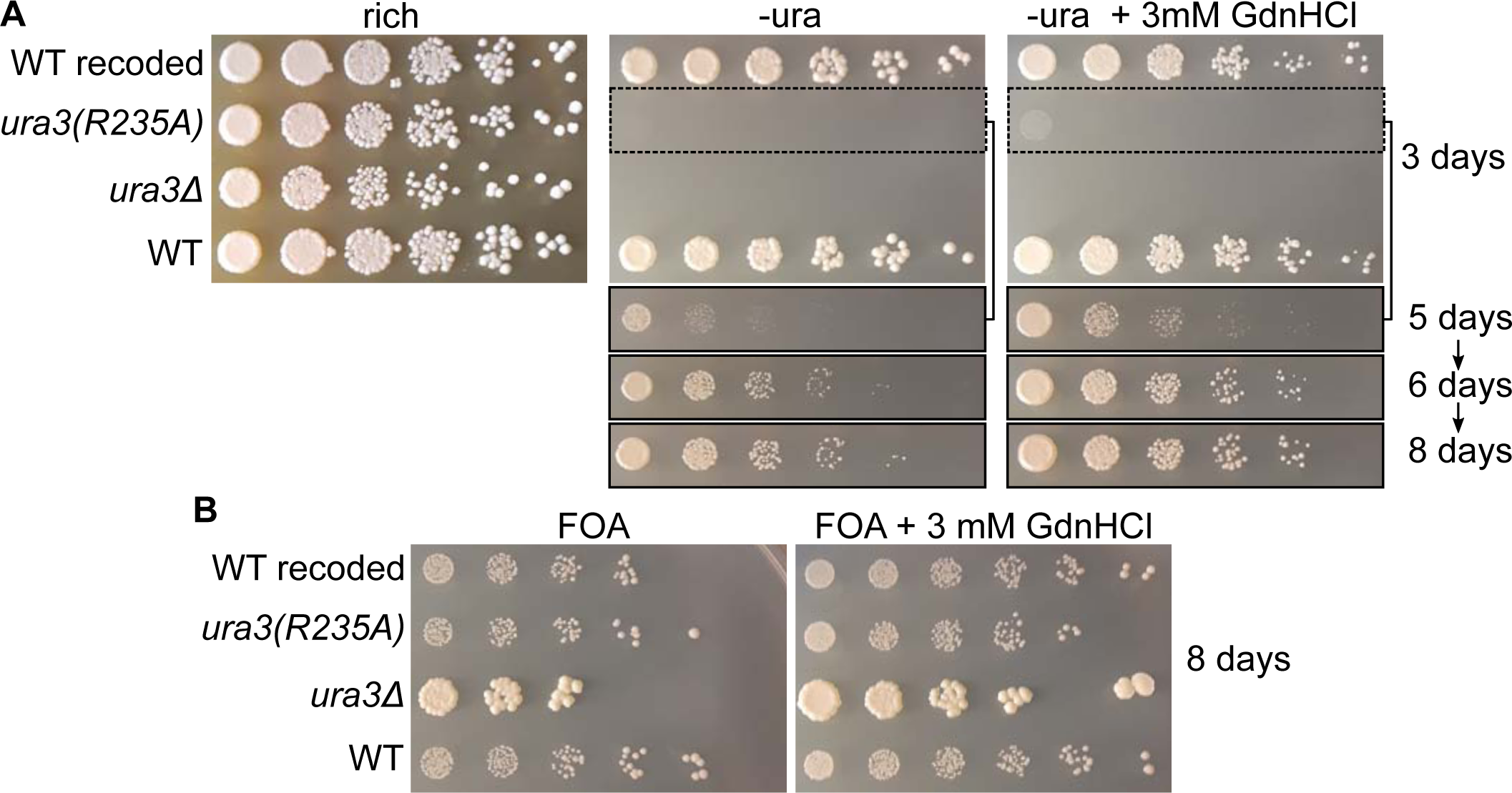
*In vivo* chemical rescue by GdnHCl of Ura3(R235A). (A) Dilution series of cells of the indicated genotypes on rich (YPD) or synthetic medium lacking uracil and thus selective for Ura3 function. Plates were incubated at 30°C for the indicated number of days prior to imaging. Strains were: “WT”, S288C; “WT recoded”, H06700; “*ura3(R253A)*”, H06701; “*ura3*Δ”, BY4741. (B) As in (A) but on medium with 20 µg/mL uracil and 400 µg/mL FOA, which is converted by Ura3 to a toxic product that inhibits colony growth.

In yeast, Ura3 converts 5-fluoro-orotic acid (FOA) to a toxic product (47). The threshold of Ura3 activity that allows growth in the absence of uracil is distinct from the threshold that inhibits colony growth on FOA (48), potentially allowing us to detect more subtle changes in Ura3 activity. On medium containing 20 µg/mL uracil and 400 µg/mL FOA, colony growth by *ura3(R235A)* cells was indistinguishable from that of cells with WT Ura3 activity, and much slower than *ura3*Δ cells (Fig. 2*B*). Addition of 3 mM GdnHCl had no discernable effect (Fig. 2*B*). We conclude from these data that GdnHCl is able to rescue the function of Ura3(R235A) in living yeast cells, but to a very minor extent.

What might explain the greater extent of *in vivo* GdnHCl rescue for ScOTC(R69G) compared to Ura3(R235A)? Each Arg residue is thought to electrostatically stabilize the substrate in the active site. The key difference may be the extent to which the Arg mutation cripples the enzyme-substrate interaction. K_M_, the substrate concentration at which a reaction rate is half-maximum, is a reliable indicator of an enzyme’s affinity for its substrate. The R57G mutation increases the K_M_ of the *in vitro* reaction catalyzed by EcOTC only about five-fold (from 50 µM to 260 µM), whereas the R235A mutation in Ura3 increases the K_M_ of the ODCase reaction over 1000-fold (1.4 µM to 1800 µM). That Ura3 relies so heavily on Arg235 to stabilize its substrate suggests that a much higher Gdm concentration would be needed for rescue *in vivo*. Indeed, the *in vitro* study estimated that restoring Ura3(R235A) activity to that of WT would require an effective Gdm concentration of 160 M (46). Our findings provide evidence that for mutant enzymes in which the mutation affects substrate binding, chemical rescue by GdnHCl *in vivo* may only be possible in cases where the mutation is only mildly destabilizing. In support of this notion, for the kinase Src the R388A substitution increased the K_M_ for ATP only about two-fold (81 µM to 182 µM) and actually decreased the K_M_ for a peptide substrate, allowing *in vivo* rescue by a non-toxic concentration of imidazole (8). Of course, if the Arg mutation has too minor of an effect on enzyme function, there may be no *in vivo* phenotype to rescue, such as what we found with β-gal(R599A) (Fig. S1); *in vitro*, the R599A mutation increased the K_M_ for the substrate *o*NPG less than two-fold (33).

### GdnHCl fails to rescue Arg-mutant asparagine synthetase

Mutation to Ala of Arg325 in *E. coli* asparagine synthetase B abolishes activity *in vitro* but 50 mM GdnHCl restores activity to ∼15% of WT (49). This enzyme has multiple substrates (ATP, glutamine, and aspartate); the R325A mutation slightly decreases the K_M_ for glutamine and increases the K_M_ for ATP just over two-fold (49). Substitution in the human homolog of the equivalent Arg (Fig. 3*A*) to His causes Asparagine Synthetase Deficiency (50). Based on these criteria, this Arg-mutant enzyme seemed like a good candidate for chemical rescue *in vivo*. The redundant *S. cerevisiae* paralogs Asn1 and Asn2 are 47% identical to *E. coli* asparagine synthetase B. Deletion of both genes renders yeast cells auxotrophic for asparagine (51). We introduced the mutation R344A, corresponding to the *E. coli* mutation (Fig. 3*A*), into Asn1 encoded on a plasmid and under control of the *GAL1/10* promoter, and looked for growth by an *asn1Δ asn2*Δ strain on media containing galactose and lacking asparagine. Only WT Asn1 allowed asparagine prototrophy, regardless of the presence or absence of 3 mM GdnHCl in the medium (Fig. 3*B*). We interpret these data as evidence that the Asn1(R344A) protein is not amenable to rescue *in vivo* by GdnHCl, at least to an extent that is detectable by our approach.

**Figure 3.**
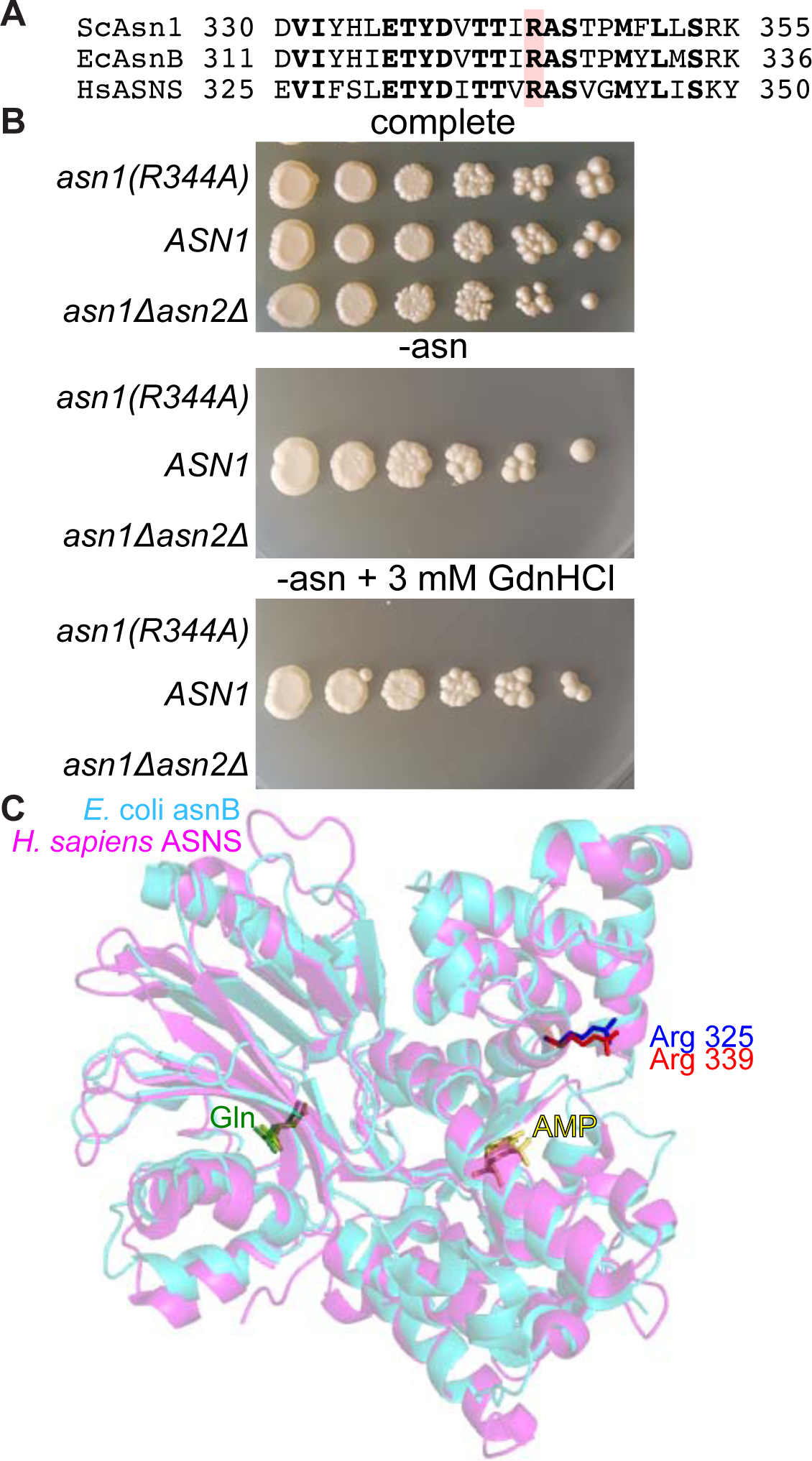
No evidence of chemical rescue of Asn1(R344A) by GdnHCl in living yeast cells. (A) Sequence alignment of a region of asparagine synthase from *S. cerevisiae* (“ScAsn1”), *E. coli* (“EcAsnB”), and *H. sapiens* (“HsASNS”). The Arg residue corresponding to EcAsnB Arg 325 is highlighted in red. (B) Dilution series of cells of the indicated genotypes on complete (but lacking uracil) or synthetic medium lacking asparagine (and uracil) and thus selective for Asn1 function (and the *URA3*-marked plasmid). Plates were incubated at 30°C for 6 days prior to imaging. Strains were: “*asn1Δasn2*Δ”, H06544 carrying empty vector plasmid pRS426; “*ASN1*”, H06544 carrying plasmid YCpU-Pgal-ASN1; “*asn1(R344A)*”, H06544 carrying plasmid YCpU-Pgal-asn1(R344A). (C) Overlay of crystal structures of EcAsnB (PDB 1CT9) and HsASNS (PDB 6GQ3) showing the locations of EcAsnB Arg 325 and HsASNS Arg 339 and two bound substrates, glutamine (“Gln”) and AMP.

A recent study also examined the R344A mutation in yeast Asn1 in the context of the filaments that Asn1 and Asn2 form in nutrient-deprived cells, the functional significance of which remain unknown (52). Asn1(R344A) was unable to form filaments (52), suggesting either that enzymatic activity is required for polymerization or that Arg 344 is required for both polymerization and activity. Structures of *E. coli* (53) and human (54) asparagine synthetase solved after the *in vitro* GdnHCl rescue study revealed that the Arg 344 equivalent in both enzymes projects away from the substrate-binding sites and towards solvent, at least in the dimeric crystal form (Fig. 3*C*). It is thus unclear how Arg in this position contributes to substrate binding or enzymatic activity, and the inter-or intramolecular contacts made by the guanidino group remain unknown. That GdnHCl rescues activity of the mutant protein *in vitro* but not *in vivo* may simply reflect a role for this Arg residue that is not recapitulated *in vitro*.

### Failure of GdnHCl to rescue a “rationally designed” mutant allele of p53

We next looked for Arg-mutant proteins that would be good candidates for *in vivo* chemical rescue by GdnHCl but had not been previously tested *in vitro*. The transcriptional activator p53 is a potent tumor suppressor and about half of all tumors express mutant forms of p53 (55). Five of the seven most common tumor-derived missense p53 mutations affect Arg residues (56). Accordingly, Arg-mutant forms of p53 have been studied extensively, including numerous attempts to rescue function using small molecules that occupy voids created by Arg substitutions (reviewed in (57)). We were particularly interested in p53(R249S) due to the identification (58) of a second-site suppressor mutation, H168R, in which the “missing” Gdm group of Arg249 is structurally replaced by the “new” Gdm group of Arg168 (59) (Fig. 4*A*). The ability of a Gdm group to occupy a void “*in trans*” despite the presence of a Ser side chain was particularly intriguing, since most previous examples of chemical rescue by GdnHCl involved Arg substitutions to Ala or Gly. We hypothesized that free Gdm might bind in the void created by the R249S substitution to restore conformational stability to the DNA-binding domain and rescue p53(R249S) function *in vivo*. Using a yeast *URA3* assay for p53 function equivalent to the one used to identify H168R as a suppressor (Fig. 4*B*), we confirmed that p53(R249S) acts as a null allele (Fig. 4*C*). However, addition of 3 mM GdnHCl to the medium had no discernable effect (Fig. 4*C*).

**Figure 4.**
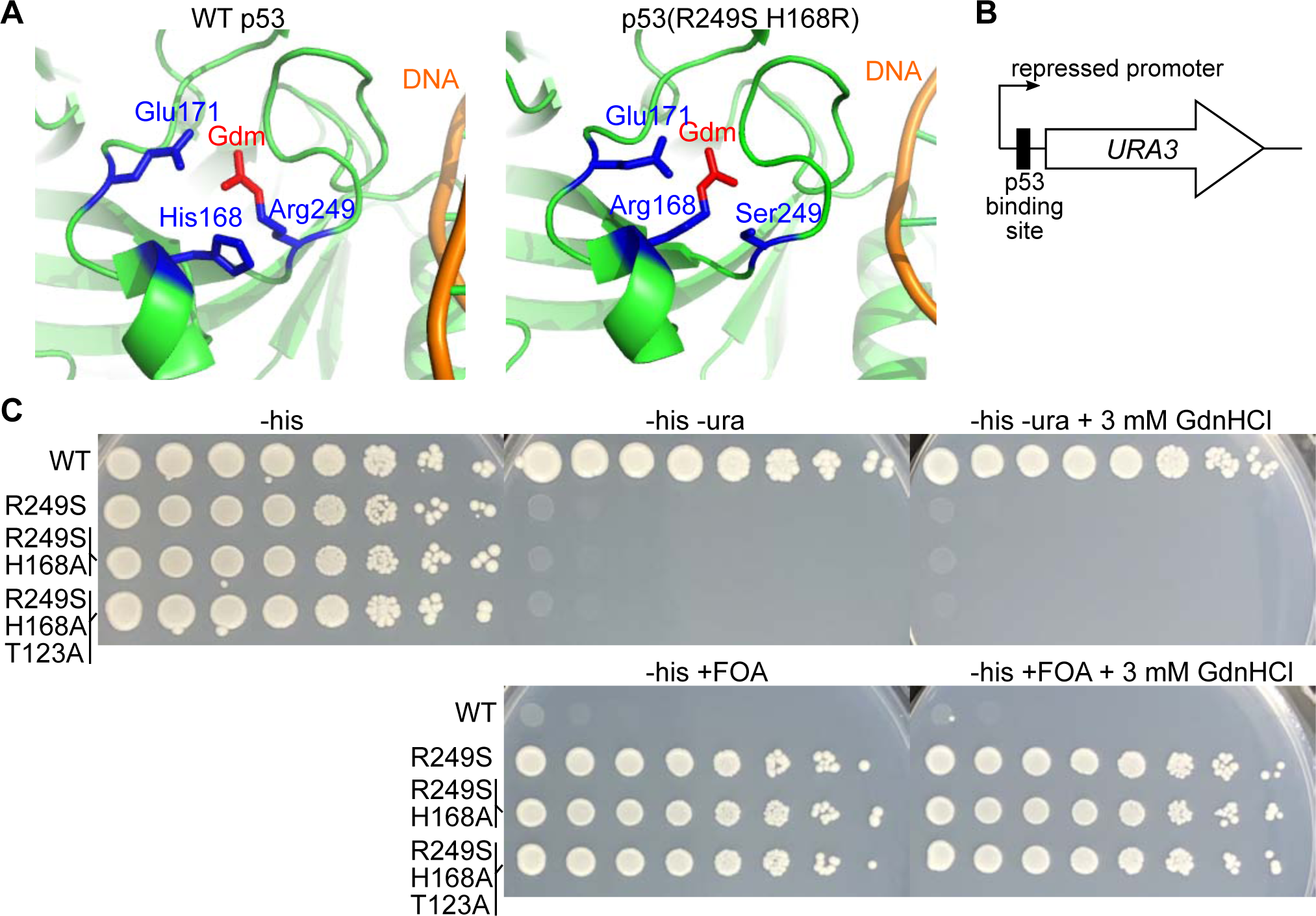
GdnHCl fails to rescue p53(R249S). (A) Crystal structures of the DNA-binding domain of WT p53 (PDB 2AC0) and p53(R249S H168R) (PDB 3D0A) showing the locations of relevant side chains and, in particular, the guanidino group (“Gdm”) occupying the same position in both proteins. Glu 171 is shown because in both cases it forms a salt bridge with the guanidino group. (B) Schematic of yeast reporter system. (C) Dilution series of cells of the reporter strain (Rby33) carrying *HIS3*-marked plasmids (“WT”, pMAM44; “R249S”, pMAM71; “R249S H168A”, pMAM70; “R249S H168A T123A”, pMAM113) were plated on synthetic medium lacking uracil (“-ura”) or containing 50 µg/mL uracil and 0.1% FOA and incubated at 30°C prior to imaging.

We suspected that the His168 side chain might interfere with occupancy of Gdm in the R249S void. However, we detected no effect of GdnHCl on the function of p53(H168A R249S) (Fig. 4*C*). Finally, we considered that H168 was isolated as a second-site suppressor of R249S only in the context of a third mutation, T123A, which makes critical but poorly understood contributions to cellular function of the triple-mutant protein (59). However, GdnHCl conferred no discernable rescue of activity to p53(T123A H168A R249S) (Fig. 4*C*). We suspect that in the case of p53(R249S) and derivatives thereof, the chemical environment surrounding the former site of Arg249 is not conducive to occupancy by free Gdm to an extent that allows the DNA-binding domain to achieve an active conformation long/often enough to confer function, at least to detection limits of our assays.

### *In vivo* chemical rescue by GdnHCl of a specific Arg-mutant actin allele

To use an unbiased approach to identify other examples of GdnHCl rescue of mutant proteins, we screened two collections of TS mutants for high-temperature colony growth in the presence and absence of 3 mM GdnHCl. One collection included multiple *cdc10* mutants, which served as convenient positive controls (Fig. 5*A*). No other mutant in this collection displayed GdnHCl rescue comparable to that of *cdc10* mutants, but one of the clearest non-*cdc10* examples was observed for *act1-105* (Fig. 5*A*,*B*). The *act1-105* allele encodes a mutant of the sole yeast actin with two surface-exposed charged residues, Glu311 and Arg312, substituted to Ala (60). The mutant cells proliferated slightly slower than WT on rich medium at room temperature (RT; ∼22°C, see Fig. S3) and not at all at 37°C unless GdnHCl was added (Fig. 5*D*). The equivalent Arg residue in human cardiac actin, Arg312, is frequently mutated in dilated cardiomyopathy (DCM), a hereditary cardiac condition (61). *act1(R312H)* mutants are TS (62) and, when synthesized *in vitro*, human cardiac actin with the R312H substitution displays prolonged interactions with chaperone proteins essential for actin folding (63). Consistent with folding defects in the mutant proteins, Arg312 makes multiple intramolecular contacts in yeast Act1 (Fig. 5*C*). Of 19 other actin mutants in the collections we screened, including six with Arg-to-Ala substitutions (Fig. 5*C*, Table S1), only *act1-105* showed any evidence of GdnHCl rescue (Fig. 5*B* and data not shown). Rescue of *act1-105* proliferation at 37°C by GdnHCl was also observed in liquid culture and was specific to GdnHCl, as we saw no effect of metformin or DMSO, two small molecules with similar chemical composition (Fig. 5*E*). These data are consistent with a model in which the Gdm ion binds at the site of the “missing” Arg at position 312 and restores intramolecular contacts that allow the mutant protein to fold, escape chaperone sequestration, and function at high temperature.

**Figure 5.**
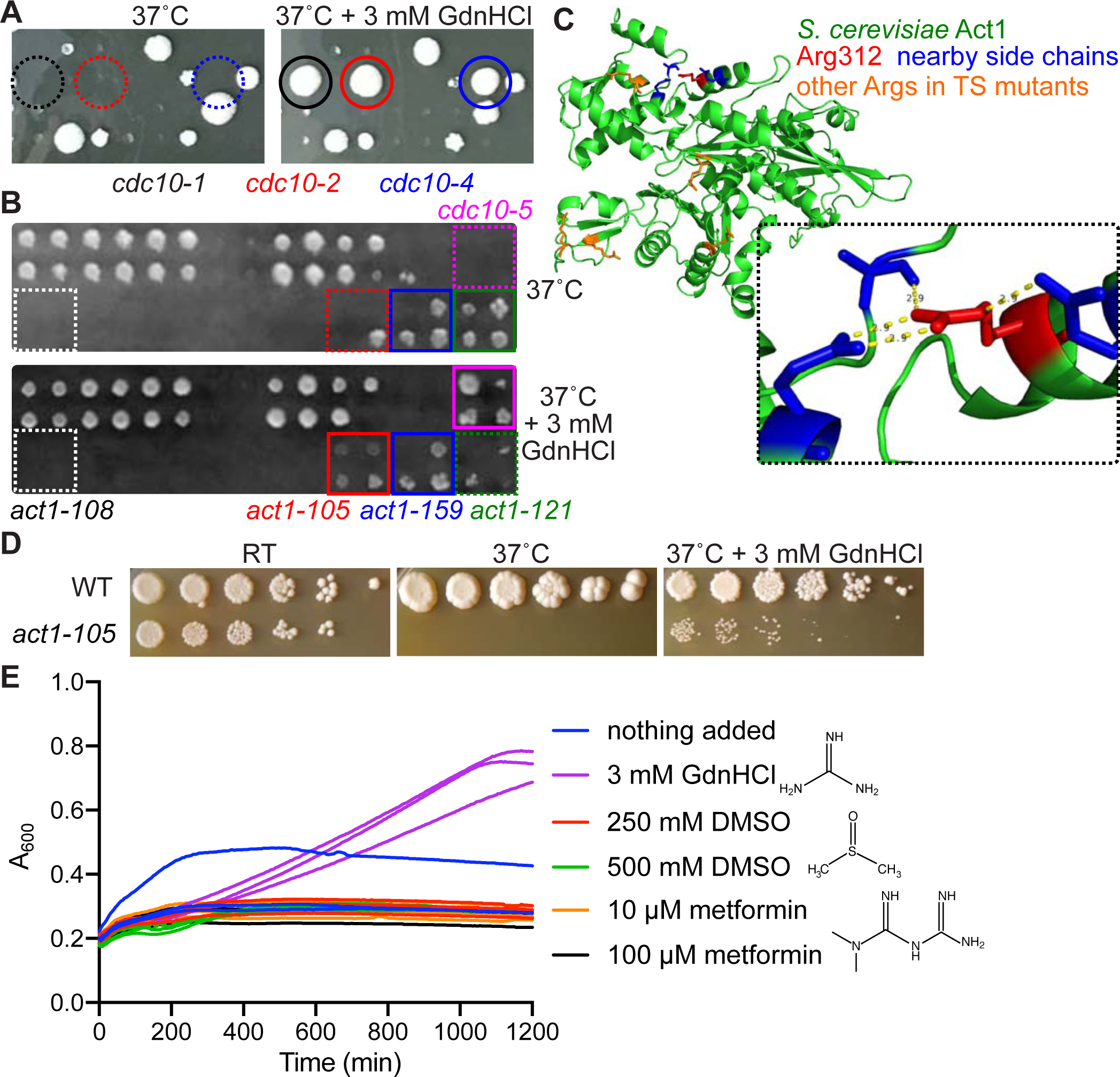
Chemical rescue by GdnHCl of Arg-mutant actin. (A) Cells from a collection of TS mutants were pinned using a screening robot to rich (YPD) plates with or without 3 mM GdnHCl and incubated at 37°C prior to imaging. Circles are color-coded with text below to indicate specific mutants. Dashed circles indicate failure to form a colony. (B) As in (A) but imaging a portion of a different plate, and using squares to indicate specific mutants. (C) Crystal structure of yeast Act1 (PDB 1YAG) showing the location of Arg 312 and nearby residues, and other Arg residues substituted in other TS mutants present in the collection we screened. Inset shows the distances (all 2.9 Å) between atoms in the guanidino group and nearby amino acids, representing presumed intramolecular contacts. (D) Dilution series of WT (BY4741) or *act1-105* cells on rich (YPD) medium at the indicated temperatures (“RT”, room temperature) with or without GdnHCl. (E) 100-µL cultures of *act1-105* cells in rich (YPD) medium with the indicated additives (three cultures for each condition) were incubated with constant shaking at 37°C in a 96-well plate and the absorbance at 600 nm was measured every 5 min.

**Table 1.** Summary of mutants rescued by small molecules

| Gene | GdnHCl<br>(3 mM) | TMAO*<br>(0.5 M) | DMSO<br>(5%) | Notes, known mutations |
| --- | --- | --- | --- | --- |
| *ACT1* | *act1-105* | *act1-111*<br>*act1-129* |  | *act1-105* (E311A, R312A)<br>*act1-111* (D222A, E224A, E226A)<br>*act1-129* (R177A, D179A) |
| *ARG3* | *arg3(R69G)* |  |  | *arg3(R69G)* not tested for TMAO, DMSO rescue |
| *ARP2* |  |  | *arp2-14* |  |
| *ARP3* |  | *arp3-D11A*<br>*arp3-G15C*<br>*arp3-H161A* | *arp3-G15C* |  |
| *CDC10* | *cdc10-1*<br>*cdc10-2*<br>*cdc10-4*<br>*cdc10-5* | *cdc10-2* |  | GdnHCl binds Cdc3, bypassing mutant Cdc10 during septin hetero-oligomerization |
| *CDC23* |  |  | *cdc23-4* | T94M |
| *ERG26* | *erg26-1* | *erg26-1* |  |  |
| *HSP10* | *hsp10-ts* |  |  | P36H |
| *KAR2* |  | *kar2-159* | *kar2-159* | G417S, variable DMSO rescue |
| *LCB1* |  | *lcb1-2*<br>*lcb1-4*<br>*lcb1-5* | *lcb1-2*<br>*lcb1-5* | *lcb1-2* (M261R)<br>*lcb1-4* (D320Y)<br>*lcb1-5* (D299Y) |
| *MCM3* | *mcm3-1* | *mcm3-1* |  | G246E |
| *ORC2* | *orc2-1* | *orc2-1*<br>*orc2-2*<br>*orc2-4* |  | *orc2-1* (P603L) |
| *PGA1* | *pga1-ts* |  |  | T25A, I28T, K40R, K98N, E147G. Not tested for TMAO, DMSO rescue. |
| *SMC3* |  | *smc3-1* | *smc3-1* |  |
| *STU1* | *stu1-5*<br>*stu1-8* |  | *stu1-5*<br>*stu1-6* | *stu1-5* (S88T Q257K T492A F483L Y534H L561S V575A T579S N596I P616L S621T)<br>*stu1-6* (mutation in N-region)<br>*stu1-8* (mutation in mid-region) |
| *TUB4* |  | *tub4-ΔDSY* | *tub4-ΔDSY* | Variable DMSO rescue |
| *URA3* | *ura3(R235A)* |  |  | *ura3(R235A)* not tested for TMAO, DMSO rescue |

\*For a complete list of TMAO rescues, see Table S2

### *In vivo* chemical rescue by GdnHCl of a mutant allele of the essential GPI-mannosyltransferase II subunit Pga1

In another TS collection we found a single, very clear example of GdnHCl rescue, for a mutant allele of the *PGA1* gene (hereafter “*pga1-ts*”) (Fig. 6*A*). Pga1 encodes a fungi-specific subunit of the essential GPI-mannosyltransferase II (64). We sequenced the *pga1-ts* allele and found five mutations predicted to result in amino acid substitutions: T25A, I28T, K40R, K98N, and E147G (Fig. 6*B*). To test if GdnHCl suppression was specific to the *pga1-ts* allele, we obtained two additional TS *pga1* alleles, *pga1-1* and *pga1-3*, each of which carries multiple distinct amino acid substitutions (Fig. 6*B*) (64). 3 mM GdnHCl failed to suppress the TS colony growth defect of *pga1-1* or *pga1-3* cells (Fig. 6*A*).

**Figure 6.**
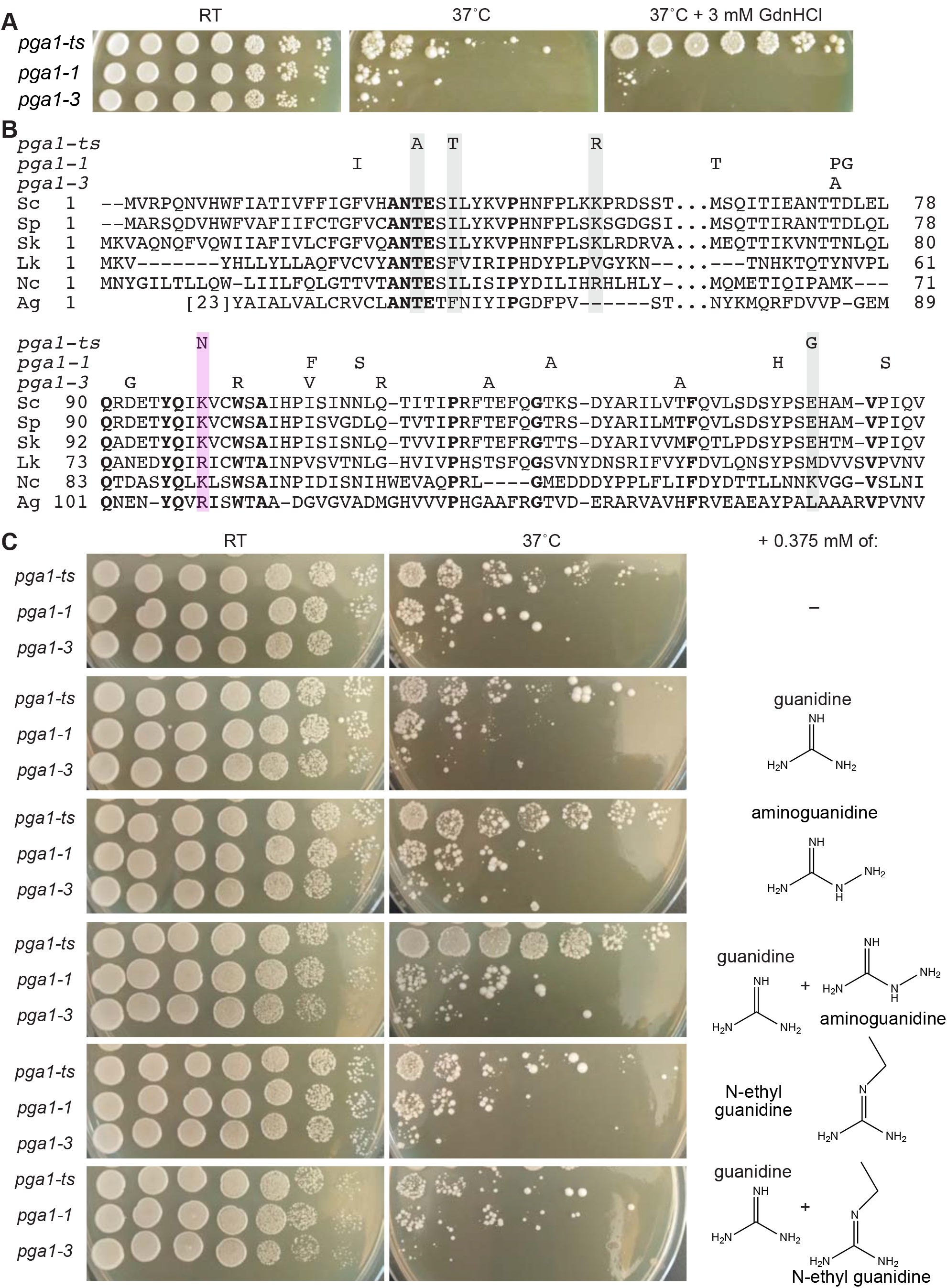
GdnHCl rescue of a Lys-mutant allele of Pga1. (A) Dilution series on rich (YPD) medium with or without 3 mM GdnHCl at the indicated temperatures for strains carrying the indicated alleles of *PGA1*. Strains were H06474, KSY182, and AKY19. (B) Sequence alignment of Pga1 sequences from various fungal species, obtained from the Yeast Genome Database and NCBI server and aligned using COBALT. Residues identical in all WT sequences are in bold. Substitutions in TS mutant ScPga1 alleles are indicated above, those in Pga1-ts are highlighted by shading. Pink, K98N substitution in Pga1-ts at a position where Arg is found in other species. Sc, *S. cerevisiae*; Sp, *S. paradoxus*; Sk, *S. kudriavzevii*; Lk, *Lachancea kluyveri*; Nc, *Naumovozyma castelli*; Ag, *Ashbya gossypii*. (C) As in (A) but with, where indicated, 0.375 mM of GdnHCl (“guanidine”) or derivatives thereof. Where multiple drugs were added, the concentration of each was 0.375 mM.

The GdnHCl derivatives aminoguanidine hydrochloride (“aGdnHCl”) and N-ethylguanidine hydrochloride (“eGdnHCl”) provide partial rescue of other Arg-mutant proteins *in vitro* in a manner that relates to the size of each molecule and thus presumably to steric compatibility with the site of binding (38). We previously showed that, relative to guanidine, aGdnHCl and eGdnHCl are less able to rescue the *cdc10* TS phenotype (19). At 0.375 mM, aGdnHCl provided a barely detectable level of rescue of *pga1-ts* growth at 37°C, while GdnHCl and eGdnHCl had no discernable effect (Fig. 6*C*). However, a cocktail of GdnHCl and aGdnHCl at a total concentration of 0.75 mM supported nearly WT colony growth by *pga1-ts* cells (Fig. 6*C*). The GdnHCl/eGdnHCl cocktail had no such effect (Fig. 6*C*). The *pga1-1* and *pga1-3* strains showed no response to any single drug or cocktail (Fig. 6*C*). These results are consistent with a model in which Gdm and, to a greater extent, aGdm make specific molecular contacts with the Pga1-ts protein that restore high-temperature function.

There is no structural information for Pga1 or any homolog, and *ab initio* structure predictions for this transmembrane protein generate low-confidence results (unpublished observations). However, only two of the substitutions in the Pga1-ts mutant protein affect highly conserved residues, and one of these, K98N, substitutes to Asn a Lys residue that is Arg in the Pga1 homologs from *Lachancea kluyveri* and *Ashbya gossypii* (Fig. 6*B*). Lys sidechains are, like those of Arg, long and positively charged at physiological pH. Thus we speculate that Gdm provides molecular contacts that are missing in the Pga1-ts protein due to the K98N substitution, and thereby restores function to the mutant cells. We note that in the crystal structure of an Arg-mutant human protein (mitochondrial aldehyde dehydrogenase R475Q) a Gdm ion occupies the position of a Gdm group not from the “missing” Arg, but from another Arg that was mispositioned due to the mutation (65). Thus in principle GdnHCl could rescue various non-Arg-mutant proteins in which a positively-charged side chain (of Arg, Lys or even His) is missing from a particular location as a direct or indirect result of a mutation.

### Rescue of a mutant allele of the C-3 sterol dehydrogenase Erg26 points to GdnHCl inhibition of specific steps in ergosterol biosynthesis

We found that the TS phenotype of the *erg26-1* mutant is partially rescued by 3 mM GdnHCl (Fig. 7*A*). The lack of 4α-carboxysterol-C3 dehydrogenase activity in *erg26-1* mutants cultured at 37°C leads to the accumulation of toxic 4-carboxysterol intermediates (Fig. 7*B*) (66). A previous search for genetic suppressors of the *erg26-1* phenotype identified mutations in *ERG1* or *ERG7* and found that these mutations decreased levels of the toxic intermediates (67). If GdnHCl inhibits the function of Erg1, Erg7, and/or other enzymes acting upstream of Erg26 in the ergosterol biosynthesis pathway, it might indirectly rescue high-temperature proliferation by *erg26-1* cells. On the other hand, for *erg* mutants with TS phenotypes that reflect a failure to produce sufficient ergosterol, rather than an accumulation of toxic intermediates, GdnHCl might inhibit proliferation at temperatures at which the mutants normally proliferate, rather than rescue proliferation at temperatures at which it otherwise fails. Looking for mutants that exhibited such GdnHCl sensitivity was complicated by the fact that at our “permissive” temperature, 3 mM GdnHCl inhibited colony growth of WT cells (Fig. 7*C*), whereas at 30°C the same concentration of GdnHCl had almost no effect (Fig. 2*A*). This effect was most pronounced when cells were diluted serially and spotted onto plates, as opposed to streaked onto plates with a toothpick (Fig. S4). Hence, it was easiest to see GdnHCl sensitivity for *erg8-1*, *erg9-ts*, *erg11-td*, and *erg20-ts*, each of which showed some degree of colony growth at 37°C in the absence of GdnHCl but none in its presence (Fig. 7*C* and see Fig. S4). The *erg10-1* mutant could not form any colony at 37°C (Fig. 7*C*), and neither *erg1* nor *erg7* was represented in these mutant collections, so we could not make conclusions about their GdnHCl sensitivity. Inhibition of any reaction in the ergosterol biosynthesis pathway upstream of Erg26, including those catalyzed by Erg8, Erg9, Erg11 and Erg20 (68–70) (Fig. 7*B*), likely reduces the production of toxic intermediates in the *erg26-1* mutant at 37°C, whereas it exacerbates ergosterol synthesis defects in the other *erg* mutants. We thus interpret our findings as evidence that GdnHCl inhibits one or more steps in ergosterol biosynthesis upstream of that catalyzed by Erg26.

**Figure 7.**
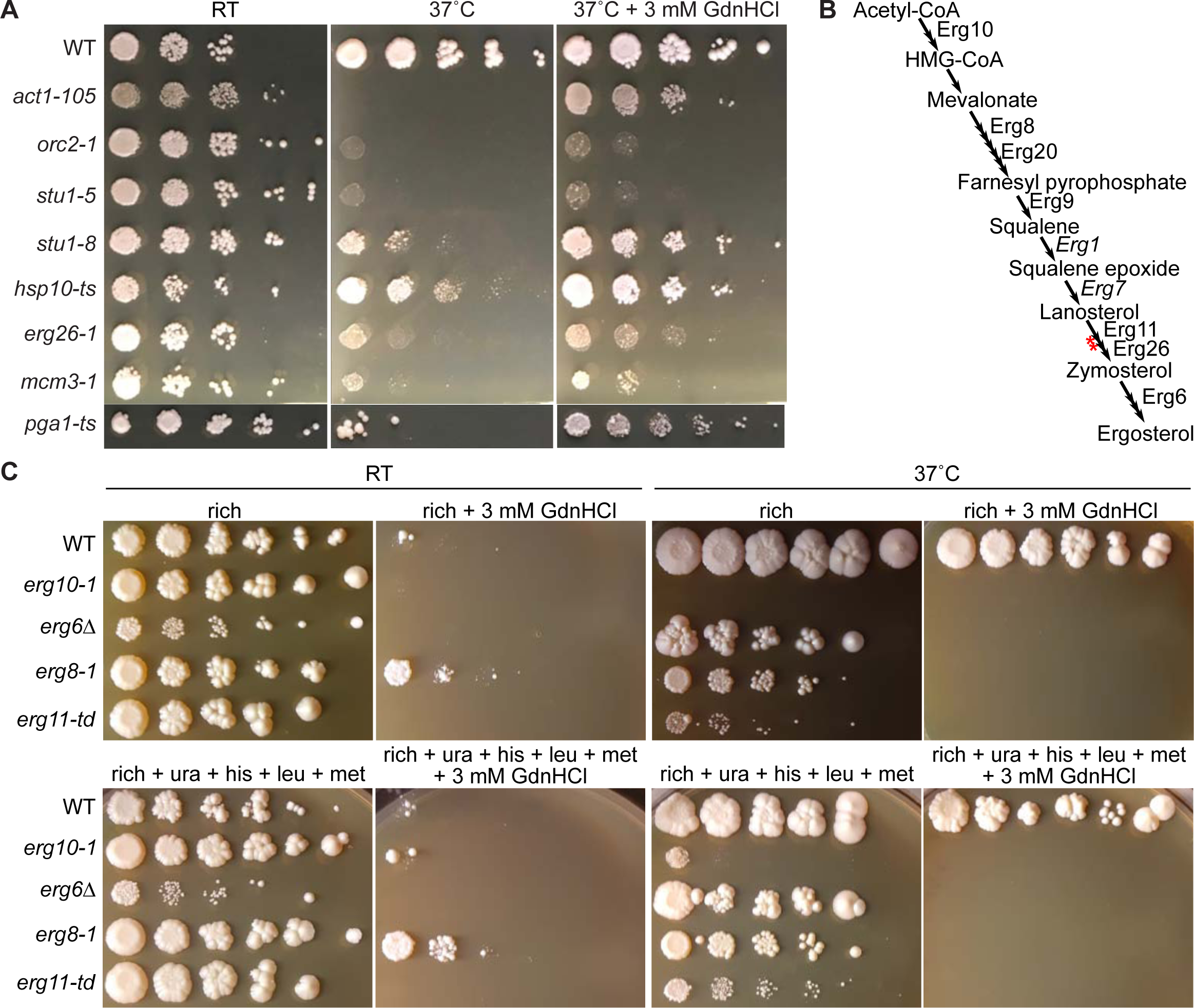
Rescue of *erg26* mutant by GdnHCl and sensitivity of WT cells and other *erg* mutants points to GdnHCl inhibition of ergosterol biosynthesis. (A) TS mutants rescued by GdnHCl identified by screening large collections. Dilution series of cells of the indicated genotypes on rich medium (YPD) with or without GdnHCl at the indicated temperatures (“RT”, room temperature). The *pga1-ts* cells were grown on a different part of the same plate as the others; the image was cropped and moved to preserve space. Strains were: “WT”, BY4741; “*act1-105*”, CBY07857; “*orc2-1*”, CBY08224; “*stu1-5*”, CBY08183; “*stu1-8*”, CBY09448; “*hsp10-ts*”, CBY10708; “*erg26-1*”, CBY11254; “*mcm3-1*”, RSY723; “*pga1-ts*”, H06474. (B) Schematic of the budding yeast ergosterol biosynthesis pathway with single reactions indicated as arrows, and labels for enzymes represented by mutants whose response to GdnHCl we tested, or, in italics, mutants known to suppress the TS defect of *erg26-1* mutants. Red asterisks indicate steps at which defects generate toxic intermediates. (C) As in (A) but with the strains of the indicated genotypes and, where indicated, rich medium was supplemented with additional nutrients for which these strains are auxotrophic (uracil, histidine, leucine and methionine). Strains were: “WT”, BY4741; “*erg10-1*”, CBY09883; “*erg6*Δ”, H06534; “*erg8-1*”, CBY08334; “*erg11-td*”, CBY04662.

We also addressed an alternative explanation for the GdnHCl sensitivity of *erg* mutants. Loss-of-function mutations in *ERG6*, the product of which acts late in the ergosterol pathway (Fig. 7*B*), increase plasma membrane fluidity and are commonly used to increase permeability of yeast cells to various small molecules (71). Indeed, we found that colony formation by *erg6*Δ mutants at 37°C was also inhibited by 3 mM GdnHCl (Fig. 7*C*). Other non-essential *erg* pathway mutants (*erg2*Δ, *erg3*Δ, and *erg4*Δ) also display increased permeability to a variety of drugs (72). If the *erg8*, *erg9*, *erg11*, or *erg20* mutant cells are similarly more permeable, they might accumulate higher intracellular concentrations of GdnHCl, mimicking the growth-inhibitory effects at temperatures ≥30°C of very high GdnHCl concentrations (≥5 mM) on WT cells (10). To test this possibility, we engineered *cdc10(D182N) erg6*Δ mutants and grew them on medium containing 1 mM GdnHCl, a concentration that provides only partial rescue of the TS phenotype of *cdc10(D182N) ERG6^+^* cells (19) and has no obvious effect on the colony growth of *CDC10*^+^ *erg6*Δ cells (Fig. S4). We predicted that if *erg* mutations generally increase permeability, then 1 mM GdnHCl should be better able to rescue the *cdc10(D182N)* TS phenotype in an *erg6*Δ background. Instead, *cdc10(D182N) erg6*Δ cells were unable to form colonies at 37°C in the absence or presence of GdnHCl (Fig. S4). Thus it is unlikely that the inhibitory effects of 3 mM GdnHCl on *erg* mutants can be explained solely by increased intracellular Gdm relative to WT.

Finally, we noticed that *erg6*Δ mutant cells exhibited a cold-sensitive colony growth defect, in that colony growth was slower than WT at RT (Fig. 7*C*) but not at 30°C (data not shown). A cold-sensitive colony growth phenotype was previously observed (at 22°C) for *erg6* mutants and was attributed to a cold-sensitive defect in tryptophan uptake (73); our *erg6*Δ mutant is prototrophic for tryptophan, but other uptake defects were not excluded in the original study. We therefore considered the possibility that in WT cells GdnHCl, like *erg6*Δ, also causes cold-sensitive defects in nutrient uptake. Interestingly, the *erg8* strain grew slightly better than WT at RT on 3 mM GdnHCl (Fig. 7*C*), providing further support for a role for membrane sterols in the inhibitory effects of GdnHCl at low temperatures. However, two pieces of evidence argue against the idea that nutrient uptake is inhibited by GdnHCl. First, while it partially improved colony growth by some *erg* mutant strains at 37°C (Fig. 7*C*), supplementing the rich medium with all nutrients for which our strains are auxotrophic did not improve growth of any strain at RT in the presence of 3 mM GdnHCl (Fig. 7*C*). Second, a fully prototrophic strain was also unable to form colonies on rich medium with 3 mM GdnHCl at RT (Fig. S4). We thus favor a model in which GdnHCl inhibits one or more specific steps in the ergosterol biosynthesis pathway. Molecular details remain unknown.

### GdnHCl rescues multiple mutants that are subject to increased proteasomal turnover

Several cases of GdnHCl rescue and sensitivity that emerged from our unbiased screens pointed to a possible inhibitory effect of GdnHCl on proteasome function. First, we found weak GdnHCl rescue of the TS growth defects of *orc2-1*, *stu1-5* and *stu1-8* mutants (Fig. 7*A*). The mutation in the *orc2-1* mutation, P603L, decreases protein expression levels ∼10-fold and this decrease together with the TS phenotype can be suppressed by mutations in subunits of the proteasome or the ubiquitin-activating enzyme Uba1 (74). The *orc2-1* allele displays positive genetic interactions with mutations in multiple genes encoding subunits of the proteasome (*PRE2*, *RPN6*, *RPN11*, *RPT5*, and *RPT6* (75)). Similarly, in *stu1-5* cells levels of the mutant protein are reduced and these levels are further reduced, to the point of synthetic lethality, when the deubiquitinylating enzyme Uba3 is mutated (76). The *stu1-5* allele also displays positive genetic interactions with mutations in many genes encoding proteasome subunits (*PUP1*, *PUP3*, *RPN1*, *RPN5*, *RPN6*, *RPN11*, *RPT3*, *RPT4*, and *RPT6* (75)). We sequenced the *stu1-5* allele and found a number of predicted amino acid substitutions, none of which alters an Arg or Lys (Table 1). The *stu1-8* mutant has not been sequenced or characterized in detail, but since it was generated in the same way as *stu1-5* and displays similar phenotypes (77), we speculate that it is also subject to proteasomal degradation at high temperature. Second, we found that a TS mutant allele of *PRE4*, which encodes a subunit of the proteasome (78), was sensitive to GdnHCl (Fig. S4). We then tested another TS proteasome mutant, *pre1-1*, and found that it was also sensitive to GdnHCl (Fig. 8*A,B*; note that the restrictive temperature for *pre1-1* is 38.5°C (79), thus colonies formed at 37°C in the absence, but not the presence, of GdnHCl). If GdnHCl inhibits proteasomal activity, this could explain the rescue of the *stu1-5* and *orc2-1* mutants and the sensitivity of *pre4* and *pre1* cells.

**Figure 8.**
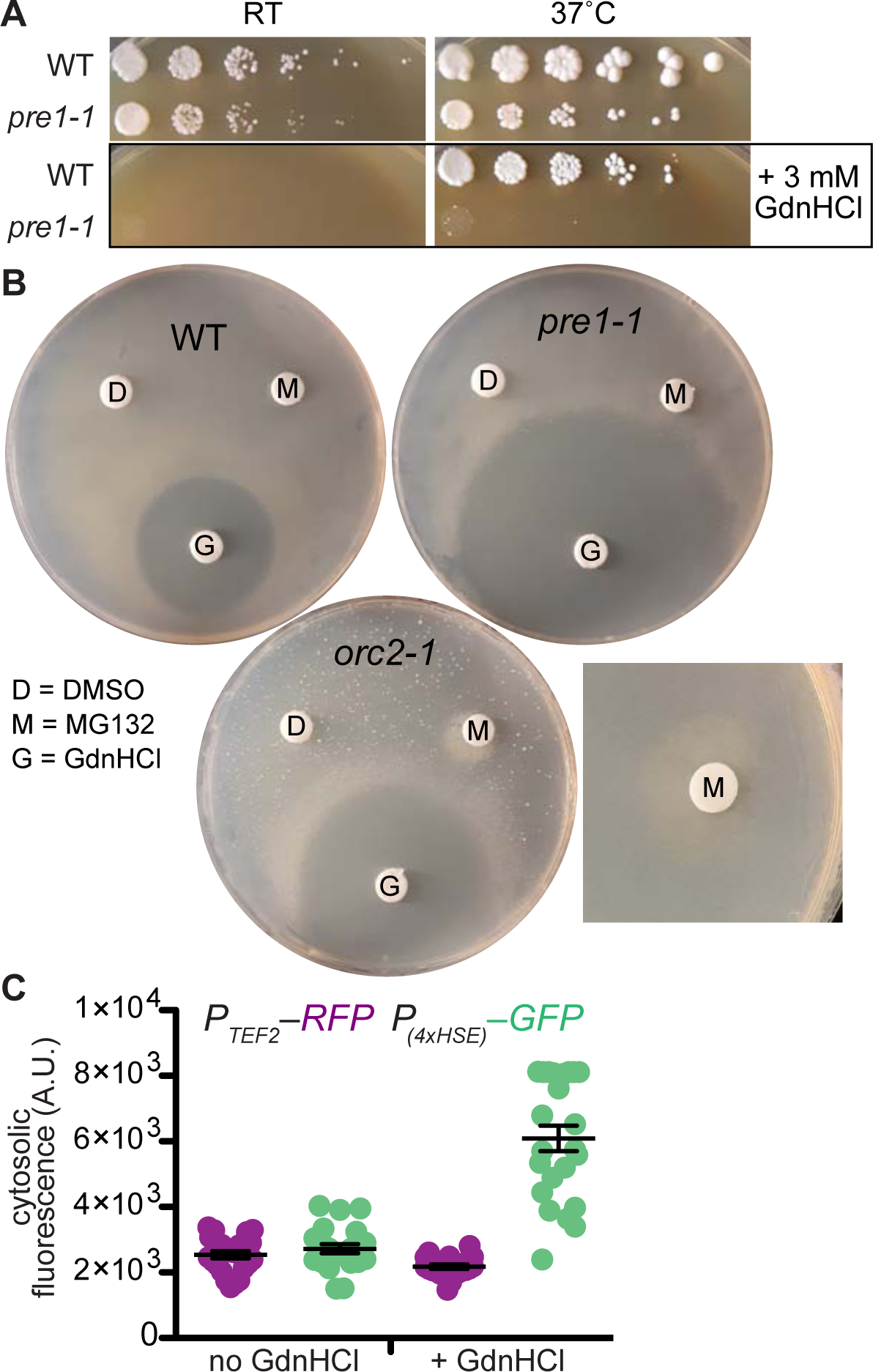
GdnHCl induces proteostatic stress and synergizes with proteasomal inhibition to rescue *orc2-1*. (A) Dilution series of BY4741 (“WT”) and CBY08187 (“*pre1-1*”) plated on rich (YPD) medium with or without 3 mM GdnHCl and incubated at room temperature or 37°C. (B) Cells from liquid rich (YPD) cultures of the indicated strains (“WT”, BY4741; “*pre1-1*”, CBY08187; “*orc2-1*”, CBY08224) were plated to medium containing Pro as the nitrogen source and 0.003% SDS to allow uptake of MG132 (156). Immediately after plating, paper discs were placed on the plate surface and 10 µL of the indicated chemicals were spotted onto the discs: 100% DMSO, 40 mM MG132 in DMSO, or 5M GdnHCl in water. Plates were then incubated at 37°C prior to imaging. The square image at right is a close-up of the MG132 disc from an independent experiment with no GdnHCl-soaked disc on the plate, ruling out any contribution from GdnHCl to the observed rescue. (C) Induction of the Hsf1-dependent cytosolic stress response measured using a strain (YJW1741) expressing RFP (mKate) from the constitutive *TEF2* promoter and GFP from an artificial Hsf1-inducible promoter containing four heat shock elements (4xHSE). Cells (23 per sample) were imaged following overnight growth at RT with or without 3 mM GdnHCl. Error bars, mean ± SEM. “A.U.”, arbitrary units.

To address this possibility, we asked whether the proteasome inhibitor MG132 also rescues the TS phenotypes of the *orc2-1* mutant. Pharmacological proteasome inhibition in *S. cerevisiae* has been previously shown to rescue the function of mutant proteins with reduced steady-state levels relative to WT (80). To include a range of MG132 concentrations, we used the paper disc assay. In addition to the expected halo of *orc2-1* rescue observed around the GdnHCl-soaked disc, we saw a zone of colony growth next to the MG132-soaked disc (Fig. 8*B*). (The sprinkling of *orc2-1* colonies dispersed across the entire plate reflects non-TS derivatives that arise spontaneously at 37°C independently of added chemicals and which we have not characterized further.)

These findings are consistent with various non-exclusive rescue mechanisms. GdnHCl might directly (or indirectly, such as by inducing chaperone expression) stabilize the conformations of mutant proteins, thereby precluding proteasomal fates. GdnHCl might directly inhibit the proteasome. GdnHCl might also indirectly inhibit the proteasome by changing the conformations of other, WT proteins in the mutant cells and increasing their targeting to the proteasome, bringing the degradation machinery near to its functional limit and allowing some molecules of the mutant protein to escape destruction. When a distinct TS allele of *pre4* is combined with hypomorphic alleles of other proteasome subunits, the double-mutant cells are hypersensitive to canavanine (81), an Arg analog that is incorporated into proteins and, like Gdm, alters protein structures and interactions. Thus our identification of *pre4* mutants as GdnHCl-hypersensitive likely reflects a mild, global proteostatic stress induced by Gdm. Indeed, at 22°C 3 mM GdnHCl clearly induced the cytosolic stress response, as assessed using a strain expressing GFP under control of an Hsf1-responsive promoter (82) (Fig. 8*B*).

By comparison, another mutant with a non-Arg substitution that we identified in our unbiased GdnHCl screens, *mcm3-1* (substitution G246E), is expressed at levels similar to WT (83), has no documented positive genetic interaction with any proteasome mutation, and is suppressed by overexpression of another subunit of the MCM complex, Mcm2 (84). These properties are most consistent with a model in which at high temperature the Mcm3(G246E) protein adopts a non-native conformation that perturbs MCM complex function, and Gdm binding to the mutant-containing complex partially restores the native conformation. [We note here that we verified that, in addition to rescuing the *mcm3-1* mutant in the isogenic TS collection we screened, GdnHCl rescued the original *mcm3-1* strain in which this mutant allele was first studied (Fig. 7*A*)]. Thus while the present data do not exclude other mechanisms, we favor a model in which Gdm restores the functions of a subset of unstable TS mutant proteins by binding directly to them (or to their direct binding partners) and altering protein conformations in ways that allow the mutant proteins to escape proteolysis. If GdnHCl directly or indirectly inhibited the proteasome, we would expect to have found a far greater number of TS mutants rescued by GdnHCl, since we suspect that proteasomal degradation at high temperature drives the TS phenotype in a significant fraction of the mutants in the collections that we screened. It is also likely that in some mutant proteins Gdm occupies molecular vacancies created by substitutions and exacerbates, rather than alleviates, mutant protein dysfunction, which may explain some of the GdnHCl-sensitive mutants that we identified.

### GdnHCl rescue or sensitivity in wild-type cells and other temperature-sensitive mutants points to a small number of cellular targets

Hsp10 is a “co-chaperonin” of Hsp60, the chaperonin that refolds numerous proteins following their import into mitochondria (85). We found weak GdnHCl rescue of the TS proliferation defect of *hsp10(P36H)* cells (Fig. 7*A*). The mutant protein shows normal thermal stability and overexpression does not improve function (86). Instead, elevated temperature appears to perturb allosteric communication between Hsp10(P36H) and Hsp60 (86). We suspect that GdnHCl could improve *hsp10(P36H)* proliferation in three non-exclusive ways. First, if there is a single essential client of Hsp60 chaperone activity that misfolds in the mutant cells at 37°C, Gdm might bind to it and directly promote acquisition of the native conformation, acting as a true “chemical chaperone”. Second, Gdm may directly act on Hsp10(P36H) and/or Hsp60 to alter the conformations of these proteins in a manner that restores allosteric communication. Third, Gdm likely inhibits Hsp78, the mitochondrial homolog of Hsp104 and of prokaryotic ClpB, which is also inhibited by GdnHCl (87). When yeast cells mutated for other mitochondrial chaperones are incubated at 37°C, mitochondria collapse into aggregates (88), and overexpression of Hsp78 partially reverses this effect (89). In *hsp10(P36H)* cells at 37°C Hsp78 may unfold mitochondrial proteins that would otherwise function normally, and GdnHCl inhibition of Hsp78 may thereby restore normal mitochondrial function. Indeed, since Hsp10 directly promotes the assembly of Hsp60 subunits into functional chaperonins (90), it may be that unassembled Hsp60 subunits themselves are targets of Hsp78 unfoldase activity. We think this third proposed mechanism is the least likely because it predicts that *hsp78*Δ should also rescue *hsp10(P36H)*, and while deletion of any one of >60 non-essential genes displays a positive genetic interaction with *hsp10(P36H)*, *hsp78*Δ (or *hsp104*Δ) is not among them (91). We note that *hsp104*Δ displays positive interactions with 10 TS mutants but not any of those we found to be rescued by GdnHCl (91), strongly suggesting that in no case is GdnHCl inhibition of Hsp104 responsible for rescue.

Inn1 is, like septins, required for cytokinesis (92, 93) and an *inn1* TS mutant displays negative genetic interactions with TS alleles of multiple septin genes (75). We found that *inn1(F59S K210E)* cells are GdnHCl sensitive at 37°C (Fig. S4). In cells with WT septins GdnHCl may slightly alter septin function in a way that is exacerbated by *inn1* mutation. Our previous studies (19) do not support the idea that GdnHCl creates a significant population of Cdc10-less hexamers in WT cells. Instead, we suspect that *inn1* sensitivity to GdnHCl reflects subtle GdnHCl-induced defects in the efficiency of septin octamer assembly at 37°C that have no overt consequence in WT cells.

We identified GdnHCl sensitivities in other TS mutants that may point to some general problem with transcription and/or mRNA processing in GdnHCl-treated cells. As shown in Fig. S4, *med6(K39R N43S M69V K75T R173G N252S D281G)*, *mpe1-ts*, *rad3(C590Y)*, and *ipa1-ts* mutants were sensitive to 3 mM GdnHCl at 37°C. Rad3 is a subunit of RNA Pol II initiation factor TFIIH (94) and Med6 is part of the RNA Pol II mediator complex (95). Mpe1 (96) is a subunit of the CPF mRNA cleavage and polyadenylation factor, which interacts physically and genetically with Ipa1 (75).

Finally, in light of the slight resistance to the inhibitory effects of 3 mM GdnHCl at RT conferred by the *erg8-1* mutation (Fig. 7*C*) we wondered if it would be possible to obtain spontaneous mutants that are resistant to inhibition by high GdnHCl concentrations. If, on the other hand, GdnHCl perturbs many distinct cellular functions, too large number of mutations would be required. We spread cells from a liquid culture of a WT haploid strain to a rich medium plate containing 10 mM GdnHCl. After 7 days incubation at 30°C, three colonies appeared, each of which was able to form new colonies when restreaked to a new 10 mM GdnHCl plate (Fig. S4). Testing for auxotrophies present in the parent strain verified that they were not contaminants (data not shown). While future work will be required to determine what change(s) promote GdnHCl tolerance in these clones, we interpret these findings as evidence that the inhibitory effects of GdnHCl in *S. cerevisiae* represent one or a small number of essential targets.

### Failure of urea to rescue any TS mutant suggests GdnHCl rescue is not due to general thermal stabilization

A model in which guanidine restores native conformations to a variety of mutant proteins, including those without Arg substitutions, is reminiscent of the non-specific conformational stabilization that has been reported for both GdnHCl and urea at subdenaturing concentrations (97). In this scenario, Gdm or urea interacts both with side chains and the backbone of the stabilized protein, suggesting that global protein stabilization is the cumulative result of many weakly stabilizing interactions, including direct electrostatic interactions and changes in surface hydration (97). In general, TS-causing mutations inhibit the ability of the mutant protein to adopt the native conformation(s) at high temperature. Hence, for some TS mutants, GdnHCl might rescue function by conferring general thermal stability. If so, we predicted that urea might do the same.

To test this hypothesis, we screened a collection of TS mutants we had previously screened for GdnHCl rescue, but this time looking for proliferation on medium containing or lacking urea at 37°C. At concentrations above 0.5 mM, urea in the medium readily enters *S. cerevisiae* cells, and enzymes that convert urea to ammonia for use as a nitrogen source are repressed if other nitrogen sources are present in the medium (98). Consequently, we predicted that in cells cultured in rich medium containing 10 mM urea, free urea should be available for interaction with intracellular proteins. However, we were unable to identify any TS mutant in this collection that was clearly rescued by, or especially sensitive to, 10 mM urea. Fig. S5 shows results of retesting several candidate mutants, each of which ultimately failed to display anything more than the slightest evidence of rescue by urea.

To test a range of urea concentrations, we chose a single TS mutant with a non-Arg substitution, *orc2-1* (substitution P603L), and placed a paper disc soaked in 5 M urea (or 5 M GdnHCl, as a positive control) on a lawn of *orc2-1* cells. Whereas a zone of proliferation was obvious surrounding the GdnHCl-soaked disc at 37°C (and a zone of no proliferation at RT), urea had no discernable effect on proliferation of the mutant cells at any temperature (Fig. S5). These negative results provide some evidence that general thermal stabilization is not sufficient to explain the rescue by GdnHCl of TS mutants, though we cannot rule out the possibility that the concentrations of available intracellular urea required for thermal stabilization of a mutant protein exceed that tested in our experiments, and may in fact be lethal.

### Chemical rescue of TS mutants by DMSO and TMAO point to effects on multiple cellular pathways, rather than on specific mutant proteins

Our findings with GdnHCl led us to investigate other naturally occurring small molecules known to alter protein conformations. DMSO is found in many natural sources including fruit (99) and marine algae (100).

In the laboratory setting DMSO is widely used as a solvent and cryoprotectant due to effects on cellular membranes, which vary with membrane lipid composition and DMSO concentration but include thinning, stiffening, softening, and probably formation of pores (101–103). DMSO also affects protein folding and stability *in vitro* (104). Like GdnHCl, DMSO (at 2.5% v/v) cures yeast cells of at least one prion (10). While screening various compounds for rescue of mutant protein function in yeast, a previous study of human cystathionine β-synthase discovered that DMSO, intended as a solvent control, itself showed an effect (27). At 614 mM, mutant function was nearly equal to WT (27). We first tested for rescue of the TS phenotype of *act1(E311A R312A)* by DMSO at 250 mM and 500 mM and saw no effect (Fig. 5*E*). We next screened a collection of TS mutants for rescue by 705 mM (5% v/v) DMSO. Colony growth by WT and most TS mutants was slower on DMSO-containing medium, particularly at low temperature (Fig. 9*A*), reminiscent of the temperature-dependent inhibition we observed with GdnHCl. Nonetheless, ten mutants representing nine genes formed colonies at 37°C only in the presence of 5% DMSO (Fig. 9*A*), indicative of chemical rescue. Based on known phenotypes and genetic interactions, we placed them into three categories with distinct possible mechanisms of DMSO action.

**Figure 9.**
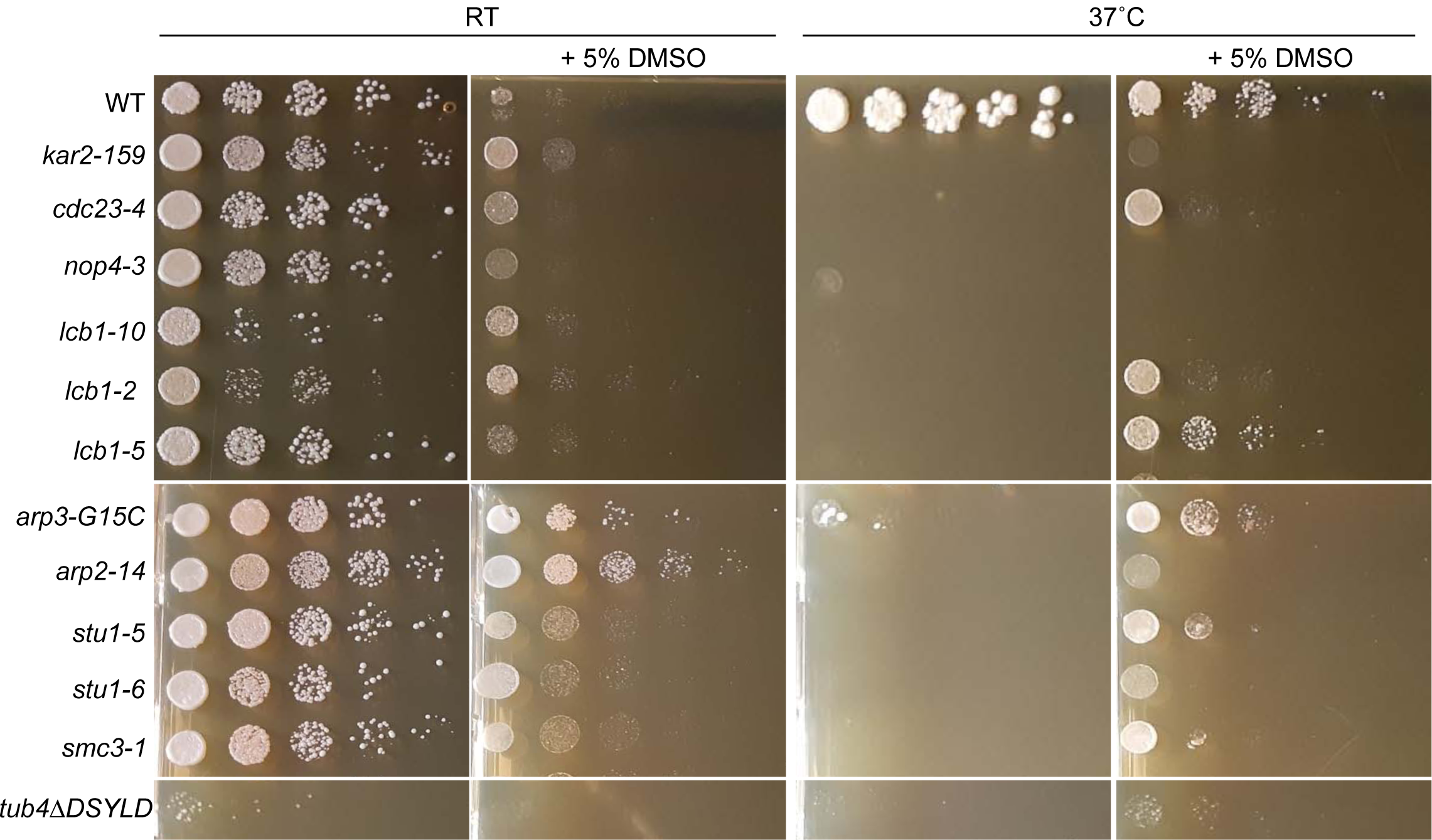
Chemical rescue of TS mutants by DMSO. Dilution series of cells of the indicated genotypes on rich medium (YPD) with or without 5% DMSO incubated at the indicated temperatures (“RT”, room temperature). Strains were: “WT”, BY4741; “*kar2-159*”, CBY07833; “*cdc23-4*”, CBY06436; “*nop4-3*”, CBY10671; “*lcb1-10*”, CBY10175; “*lcb1-2*”, CBY10161; “*lcb1-5*”, CBY10156; “*arp3-G15C*”, CBY08235; “*arp2-14*”, CBY04958; “*stu1-5*”, CBY08183; “*stu1-6*”, CBY10325; “*smc3-1*”, CBY08012; “*tub4ΔDSYLD*”, CBY11146.

DMSO weakly rescued *kar2-159* (Fig. 9*A*), a TS allele (substitution G417S) of the Hsp70-family ER chaperone Kar2, the yeast homolog of mammalian BiP (Binding immunoglobulin Protein) (105). Gene expression analysis of *S. cerevisiae* cells exposed to 1M DMSO found that Kar2 transcript levels increase slightly (106). Hence a trivial explanation is that elevated levels of the Kar2(G417S) protein compensate for its dysfunction. Alternatively, if DMSO generally improves protein folding, it might improve the folding of essential proteins otherwise prone to misfolding when Kar2 is non-functional. If so, we predicted that DMSO might also rescue a TS mutant in which Hsp70 function in folding cytosolic proteins fails at 37°C. Due to redundancy between multiple cytosolic Hsp70 family members, no cytosolic Hsp70 is represented in the collection of single-gene TS mutants that we screened. Instead, we tested two strains, one in which three of the four *SSA* (Stress-Seventy subfamily A) genes is deleted and only WT *SSA1* remains, and an otherwise isogenic strain carrying a TS allele of *SSA1* (107). 5% DMSO failed to rescue the *ssa1-ts* strain and blocked colony growth by the *SSA1* strain at 37°C (Fig. S6). These results are consistent with a model in which DMSO alters protein conformations, but in ways that do not generally substitute for the action of Hsp70 chaperones, and may in fact exacerbate defects in Hsp70-mutant cells, depending on the affected cellular compartment.

Several mutants rescued by DMSO represent proteins directly or indirectly involved in lipid synthesis and membrane dynamics. Lcb1 (*lcb1-2* and *lcb1-5*, Fig. 9*A*) is a serine palmitoyltransferase required for sphingolipid synthesis (108). The substitutions in these alleles are distinct (Table 1) and other *lcb1* mutants in the collection were not rescued (Fig. 9*A* and data not shown), suggesting that DMSO does not specifically act on one mutant form of the protein and also that DMSO does not prevent or counteract the effects of Lcb1 dysfunction in general. Addition of DMSO to *S. cerevisiae* is known to upregulate lipid synthesis genes, resulting in increased phospholipid content in membranes, which presumably counteracts direct membrane effects of the drug (109). Accordingly, in one model for DMSO rescue of *lcb1* mutants, proliferation fails due to altered membrane properties of the mutant cells cultured at 37°C, resulting from altered sphingolipid content, and DMSO restores membrane properties to something compatible with proliferation. However, sphingoid bases also act as signaling molecules and thereby influence many cellular processes, including the response to high temperature itself: in cells carrying a distinct TS allele of *LCB1* that was not present in the collection we screened, proliferation at 37°C is restored by overexpressing ubiquitin, which appears to promote proteasomal clearance of misfolded proteins without restoring normal sphingolipid content (110). Thus DMSO could rescue *lcb1* mutants in a number of ways among which our current data are unable to distinguish.

Arp2 and Arp3 are 30% identical, actin-related proteins that are essential subunits of a heptameric complex that nucleates actin filaments (111) and is required for the motility of cortical actin patches (112) and, in turn, the internalization step of endocytosis (112). While *arp2-14* is the only allele of *ARP2* in the collection we screened, *arp3-G15C* was the only of six *arp3* alleles to be scored as rescued by DMSO in our screen (Table 1, Table S1). It is unclear why DMSO rescue was specific to *arp3-G15C*, though the study that characterized these *arp3* mutants found a unique combination of phenotypes for *arp3-G15C* with regard to sensitivity to temperature, 0.9 M NaCl, and 3% formamide (113). In cultured mammalian cells, DMSO and other amphiphilic compounds alter membrane tension and thereby increase the rate of endocytosis (114).

Hence in a simple model for DMSO rescue of Arp2/3 mutants, altered membrane tension facilitates endocytic events that are otherwise stalled at the plasma membrane. Interestingly, *arp2-14* and *arp3-G15C* cells also proliferated better than WT at RT in the presence of 5% DMSO (Fig. 9*A*). Thus DMSO may synergize with low temperature to alter membrane properties in a way that perturbs endocytosis and is counteracted by subtle changes in actin patch motility in these *arp* mutants. A previous screen of the *S. cerevisiae* homozygous diploid deletion collection identified nine mutants that were more resistant than WT to 8% DMSO (115), none of which has any obvious functional relationship to endocytosis or membrane tension. However, a recent study found that *chm7*Δ also confers improved colony growth in the presence of 4% DMSO, which presumably relates to Chm7’s function in membrane dynamics in the ER and nuclear envelope (116). Why loss of Chm7 would ameliorate effects of DMSO on membrane properties remains unclear.

The final category of TS mutants rescued by DMSO – *stu1-5, stu1-6, tub4ΔDSYLD*, *cdc23-4*, and *smc3-1* – had links to microtubule function, which in yeast is mostly restricted to the mitotic spindle. Stu1 directly binds microtubules and mutations in tubulin subunits suppress the TS defect of *stu1-5* mutants (77). *TUB4* encodes gamma tubulin, which nucleates microtubules, and drugs that alter the stability of microtubules rescue the TS phenotype of the *tub4ΔDSYLD* mutant (117). [We note that DMSO rescue of *tub4ΔDSYLD* was subtle at best and altogether absent in some experiments (see Fig. S7).] Cdc23 is a subunit of the anaphase-promoting complex that localizes to microtubule ends (118). Smc3 is a subunit of the cohesin complex that holds sister chromatids together, contributing to tension that is crucial for stable microtubule– kinetochore attachments (119). A recent study (120) identified a genetic interaction between *smc3-1* and *irc15*Δ, which eliminates a microtubule-binding protein that regulates microtubule dynamics (121). DMSO exerts profound effects on purified tubulin *in vitro*, lowering the critical concentration for microtubule polymerization (122) and lifting the requirement for microtubule-associated proteins (123). In cultured mouse oocytes, exposure to ∼10% DMSO rapidly induced the formation of cold-resistant microtubules (124). We therefore speculate that direct DMSO effects on microtubule stability/dynamics underlie chemical rescue of this category of mutants.

Studies with mutant alleles of human cystathionine β-synthase expressed in yeast found that some mutant proteins could be rescued by DMSO or TMAO, whereas other mutants could be rescued only by TMAO (27). We screened the same collection of TS mutants with 0.5 M TMAO at 37°C and found nearly 140 candidate examples of rescue, six of which were also rescued by DMSO (Table 1, Table S2, Fig. S7, Fig. S8). Put another way, DMSO rescued <5% of the mutants rescued by TMAO, but TMAO rescued over half of the mutants that were rescued by DMSO. Validating all the candidates by dilution series would have been impractical, so we tested only a subset (Fig. 10 and Fig. S7). Four alleles of *PKC1*, which encodes a kinase crucial for cell wall integrity, are known to be rescued by a variety of osmolytes (125, 126); higher osmolarity of the medium counteracts intracellular turgor pressure and prevents cell lysis when cell walls are structurally compromised. Each of these *pck1* mutants was also rescued by TMAO (Fig. 10*A*). One allele of *cdc10* showed weak rescue by TMAO (Table 1 and see Fig. 10*C*). Septins are also required for proper cell wall biogenesis (127, 128), and two other septin-mutant strains, *cdc12-1* and *cdc12-6*, are also known to be partially rescued by the osmolyte sorbitol (129). Since and all TS septin mutants show similar cellular defects at the restrictive temperature (130), we suspected that TMAO rescue of *cdc10-2* may also represent a case of “osmotic support”. Indeed, at 37°C 1 M sorbitol rescued *pkc1-1* and *cdc10-2* (Fig. 10*B*). However, it was unclear why the *cdc12-1* mutant and other *cdc10* mutants in the collection we screened did not show TMAO rescue. Sorbitol rescue of *cdc12-1* in the published study was only apparent up to 34°C (129), and we confirmed that at 37°C 1 M sorbitol failed to rescue the *cdc12-1* mutant present in the collection (Fig. 10*B*). We assessed a range of temperatures in liquid cultures and found that *cdc10-2* has a higher permissive temperature than *cdc12-1* and the other *cdc10* mutants in the collection (Fig. S7). Thus only for *cdc10-2* did moderate TMAO rescue exceed the threshold of detection in our screen. We conclude that for many mutants TMAO acts as a general osmolyte to prevent cell lysis when the integrity of the cell wall is compromised.

**Figure 10.**
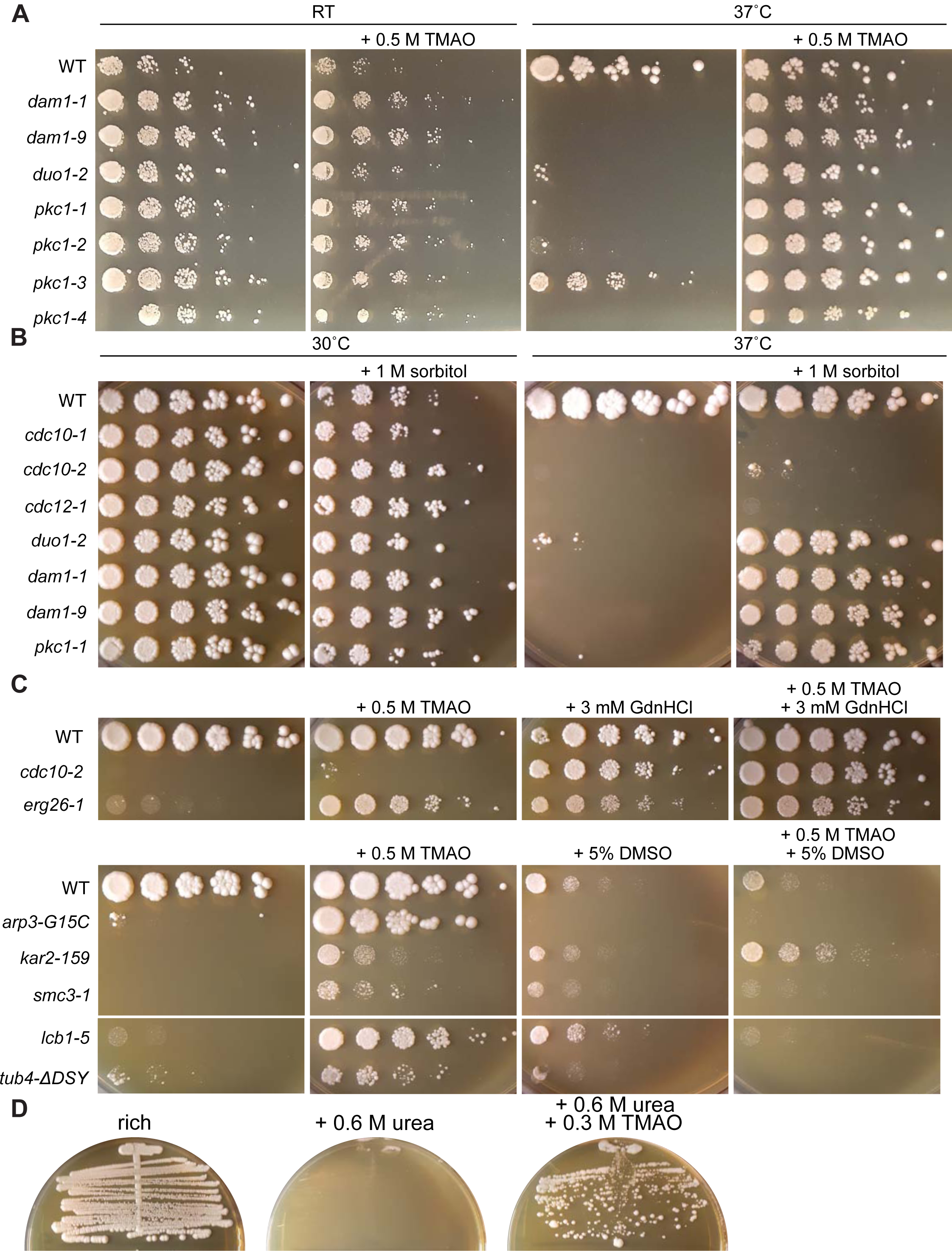
Chemical rescue of TS mutants by TMAO. (A) Dilution series of cells of the indicated genotypes on rich medium (YPD) with or without 0.5 M TMAO incubated at the indicated temperatures (“RT”, room temperature). Strains were: “WT”, BY4741; “*dam1-1*”, CBY07974; “*dam1-9*”, CBY07966; “*duo1-2*”, CBY07958; “*pkc1-1*”, CBY09957; “*pkc1-2*”, CBY09962; “*pck1-3*”, CBY09972; “*pkc1-4*”, CBY09967. (B) As in (A) but with different mutants and 1 M sorbitol instead of TMAO. Strains were: “WT”, BY4741; “*cdc10-1*”, CBY06417; “*cdc10-2*”, CBY06420; “*cdc12-1*”, CBY05110; “*duo1-2*”, CBY07958; “*dam1-1*”, CBY07974; “*dam1-9*”, CBY07966; “*pkc1-1*”, CBY09957. (C) As in (A) but with different mutants, 0.5 M TMAO, 3 mM GdnHCl and/or 5% DMSO, and all plates were incubated at 37°C. Strains were: “WT”, BY4741; “*cdc10-2*”, CBY06420; “*arp3-G15C*”, CBY08235; “*kar2-159*”, CBY07833; “*smc3-1*”, CBY08012; “*lcb1-5*”, CBY10156; “*tub4ΔDSYLD*”, CBY11146. (D) BY4741 was streaked on rich (YPD) medium with or without 600 mM urea or 300 mM TMAO and incubated at 37°C for 4 days prior to imaging.

Might we have found even more examples of TMAO rescue had we screened at lower temperatures? Comparison of the annotated restrictive temperatures of the mutant collection as a whole with those that showed apparent TMAO rescue at 37°C suggested a slight bias towards mutants with restrictive temperatures closer to 37°C, but not a strict threshold below which no rescue is possible (Fig. S7). Thus while it appears to act quite generally, TMAO does not universally improve mutant protein function; indeed, some mutants were inhibited by TMAO (see Table S2, Fig. S8 and the *brl1-2221* and *brl1-3231* mutants in Fig. S7).

Dam1 and Duo1 are subunits of the Dam1 complex (also called DASH) that links kinetochores to the microtubules of the mitotic spindle (131, 132). TMAO rescued *duo1-2* and multiple distinct TS alleles of *DAM1* (Fig. 10*A*, Table 1, Table S2), and also rescued *tub4ΔDSLYD* (Fig. 10*C*). Multiple studies in the literature observed rescue of TS mutants with spindle defects by sorbitol (133, 134), and sorbitol provides resistance to WT cells against drugs that destabilize microtubules, leading to the idea that osmotic support stabilizes microtubules (134). We found that, like TMAO (Fig. 10*A*), 1 M sorbitol also rescued *dam1-1*, *dam1-9*, and *duo1-2* (Fig. 10*B*). We propose that in these cases TMAO acts as an osmolyte to stabilize microtubules.

It is worth noting that a recent study found that upon exposure to elevated temperatures yeast cells accumulate intracellular trehalose as a way to increase cytosolic viscosity and maintain normal kinetics of protein folding, higher-order assembly, and enzymatic reactions in the face of increased kinetic energy (135). Thus at 37°C a mutant protein faces not only an intrinsically different folding environment than at RT, but once folded any subsequent intermolecular interactions, including between enzyme and substrate, are also fundamentally altered by the presence of osmolytes like trehalose and TMAO. TS mutant rescue by TMAO could result from multiple direct and indirect effects, which may explain why TMAO was able to rescue so many different TS mutants.

The human cystathionine β-synthase studies found that a cocktail including TMAO, DMSO, and sorbitol appeared to have a synergistic rescue effect (27). We tested cocktails of TMAO plus GdnHCl and TMAO plus DMSO on selected mutants and found that in some cases the combinations provided approximately additive positive effects on mutant proliferation (*e.g.* TMAO plus GdnHCl for *cdc10-2*) whereas in others (*e.g.* TMAO plus DMSO for *lcb1-5*) the combination eliminated rescue altogether (Fig. 10*C*). We imagine that additive effects could arise from a combination of a direct effect of one small molecule on the mutant protein plus a direct effect of the other small molecule on a cellular process altered as a result of the mutation. For example, Gdm binding to the septin Cdc3 bypasses incorporation of the mutant Cdc10 (19), but the resulting hexameric septin complexes are not fully functional, leading at 37°C to slight cell wall/cytokinesis defects and slower colony growth that are further remedied by the osmotic support TMAO provides. On the other hand, in *lcb1-5* cells TMAO might change the conformation of the Lcb1(D299Y) protein or its reaction kinetics in a way that increases serine palmitoyltransferase activity, but the resulting lipid composition may be incompatible with the direct membrane effects of DMSO.

Finally, given the apparent effectiveness of TMAO in rescuing TS mutant protein function, and the natural accumulation of TMAO in organisms with high intracellular urea, we predicted that if the toxic effects of high concentrations of exogenous urea on *S. cerevisiae* reflect mis- or un-folding of essential proteins, then TMAO might counteract these effects and restore viability. To test this prediction, we streaked WT haploid cells on rich medium with 600 mM urea in the absence or presence of 300 mM TMAO, a concentration chosen to mimic the 2:1 urea:TMAO ratio found in many marine organisms (24). Urea completely blocked colony formation at 37°C and the addition of TMAO almost completely reversed this effect (Fig. 10*D*). While there are many possible mechanistic explanations for this result, it is consistent with a model in which exogenous urea denatures proteins and TMAO counteracts this effect, as in the organisms inside which both molecules are found naturally.

### Conclusions

Our unbiased screens of large collections of mutants and our targeted tests of Arg-mutant proteins illustrate just how lucky we were to identify GdnHCl rescue of *cdc10* mutants (19). Only one other mutant, *pga1-ts*, showed a similar extent of GdnHCl rescue. Weaker GdnHCl rescue of *act1(E311A R312A)* but not ∼30 other Arg-to-small-side-chain mutants in the same TS collection (Table S1) confirms that such examples are rare and cannot be predicted based on structure alone. Our findings further suggest that GdnHCl rescue is most likely to be effective when Gdm acts as a “chemical chaperone” to promote native folding by restoring specific intramolecular contacts, rather than replacing in an enzyme the functional group of a missing Arg residue that makes direct contact with a substrate. Towards the latter goal, modified compounds able to covalently attach to the mutant protein (*i.e.*, “chemical repair”) may be more effective.

Nonetheless, in the context of human disease, even the slightest increase in function might improve quality of life. Due to the abundance of mutation-prone CpG dinucleotides in Arg-encoding codons, Arg is the most frequently mutated residue in human disease (136). To estimate just how many human diseases arise from Arg mutations, we analyzed a published list of single missense mutations linked to human disease and found that 18% (538 of 2960) represent Arg substitutions (137). If GdnHCl has the potential to restore function to even a fraction of these, then it should be considered as a possible therapeutic. With regard to side effects, our findings also provide additional insights into the cellular processes likely to be affected directly by GdnHCl in WT cells (*e.g.* ergosterol/cholesterol synthesis) and conditions in which these effects are most pronounced (low temperature). Our results with other small molecules suggest that, unlike them, GdnHCl does not act very generally to stabilize protein conformations *in vivo*. DMSO and TMAO likely have their own direct effects on specific cellular processes but each, particularly TMAO, also appears to act on protein conformations in general ways that ameliorate functional defects in multiple mutant proteins. DMSO is found in many natural foods (99) and has a long history of over-the-counter use as a remedy for many ailments, whereas TMAO is produced by gut microbes and enters the circulatory system, particularly following high-protein meals (138). Hence, these two molecules also have the potential to influence cellular function in humans, whether administered intentionally or otherwise.

In the laboratory context, our findings provide multiple clear illustrations of how standard *in vitro* conditions can fail to recapitulate conditions encountered by proteins in living cells, and also establish potential new tools in the form of “conditional alleles” of numerous genes whose activity can be controlled pharmacologically. Apart from these laboratory research and clinical therapeutic considerations, by modifying the chemical environment in which proteins fold and function and potentially buffering fitness against the effects of mutations, these naturally occurring small molecules have almost certainly influenced the evolution of protein sequence space in at least some organisms in some environments.

## EXPERIMENTAL PROCEDURES

### Microbial strains and plasmids

We screened two collections of TS mutants. Screening methods are provided below. The “Hieter collection” has the following genotype for each haploid TS mutant: *MAT***a** *ura3Δ0 leu2Δ0 his3Δ1 can1Δ::LEU2-PMFA1::HIS3 yfeg-ts::URA3*, where “*yfeg-ts*” is “your favorite essential gene”, of which 600 are represented (139). The allele at *MET15* was either *met15Δ0* or *MET15*^+^, and at *LYS2* was either *lys2Δ0* or *LYS2^+^*, but this information is not available for all mutants. From this collection, a separate stock of the *pga1-ts* mutant was made and assigned the inventory number H06474. The “Boone collection” that we screened comprises 684 mutants representing 444 essential genes, based on the parent strain BY4741 (*MAT***a** *his3Δ1 leu2Δ0 met15Δ0 ura3Δ0*) with each TS allele integrated using a *kanMX* marker 3’ of the stop codon (140). Table S1 is an updated copy of a previously published table from the Boone lab (140) that lists the mutants in the version of the collection we screened along with the temperature(s) at which the TS phenotype is manifested and, where known, the mutations in the TS allele. The following are yeast strains not in these collections. S288C is *MATα SUC2 ho mal mel gal2 CUP1 flo1 flo8-1 hap1* (141). BY4741 is derived from S288C and is *MAT***a** *his3Δ1 leu2Δ0 met15Δ0 ura3Δ0* (142). The *erg6Δ::kanMX* derivative from the BY4741 deletion collection (143, 144) is named H06534 in our lab collection and was confirmed by cycloheximide sensitivity (145). The *cdc10(D182N) erg6*Δ strain H06537 was made by crossing H06534 with the *cdc10(D182N)* strain DDY2334, a gift from David Drubin (University of California Berkeley), sporulating the resulting diploid, and dissecting tetrads. Other strains from the deletion collections were confirmed by other phenotypes: the *arg3Δ::kanMX* derivative of BY4741 (strain H06538) was confirmed by arginine auxotrophy, the *asn1Δ::kanMX* derivative of BY4741 and *asn2Δ::kanMX* derivative of BY4742 (*MATα his3Δ1 leu2Δ0 lys2Δ0 ura3Δ0*, (142)) were confirmed by the asparagine auxotrophy of H06544, the haploid *asn1Δ::kanMX asn2Δ::kanMX* haploid strain created by mating those two mutants, sporulating the resulting diploid, and dissecting tetrads. Strains H06744, H06745 and H06746 are spontaneous derivatives of BY4741 derived by plating to 10 mM GdnHCl. Strains AKY19 (*pga1-1*) and KSY182 (*pga1-3*) are *MET15* derivatives of BY4741 and have *LEU2* integrated at the *PGA1* locus (64). The p53 reporter strain RBy33 is *MAT***a** *his3Δ200 leu2Δ1 lys2Δ202 trp1Δ63 1cUAS53::URA3* (146). The cytosolic stress reporter strain YJW1741 is *MATα his3Δ1 leu2Δ0 cyh2 can1Δ::PSTE2-spHIS5 lyp1Δ::PSTE3-LEU2 ura3Δ::PTEF2-mKate–URA3–4xHSE-EmeraldGFP* (147). The original *mcm3-1* mutant strain Rc61-3c is *MATα his4Δ34 leu2-3,112 ura3-52 mcm3(G246E)* (83). The *ssa*Δ strains ECY487 (*SSA1*) and ECY567 (*ssa1-45BKD*) are both *MATα leu2-3,112 his3-11,3-15 trp1Δ1 ura3-52 ssa2::LEU2 ssa3::TRP1 ssa4::LYS2* (43).

The *E. coli* strains used were DH5*α* (*fhuA2 lac(del)U169 phoA glnV44 Φ80’ lacZ(del)M15 gyrA96 recA1 relA1 endA1 thi-1 hsdR17*) and RS302 (alias AMA1004), which is based on strain M182 (148) with the following modifications: *hsdR-del(ara, leu) trpC9830 leuB6 ara+ Δ(laclPOZ)C29 lacY^+^*.

All yeast transformations were performed using the Frozen-EZ Yeast Transformation II Kit (#T2001, Zymo Research). To integrate the *ura3(R235A)* allele, the Cas9-encoding plasmid (Cas9-NAT (149), a gift from Yong-Su Jin, Addgene plasmid # 64329; http://n2t.net/addgene:64329; RRID:Addgene_64329) was transformed into S288C and selected with nourseothricin. A geneblock repair template (see Fig. S2) containing the R235A mutation or a recoded version of WT *URA3* was amplified by PCR using primers ura3_GBfw (5’TCCATGGAGGGCACAGTTAA) and ura3_GBre (5’CACCCTCTACCTTAGCATCC) and the PCR product was co-transformed into S288C cells carrying the Cas9 plasmid with a gRNA-encoding plasmid, gRNA-ura-HYB (149) (a gift from Yong-Su Jin, Addgene plasmid #64330; http://n2t.net/addgene:64330; RRID:Addgene_64330). Transformants arising on YPD medium with nourseothricin and hygromycin B were screened by isolation of genomic DNA, PCR of the *URA3* coding sequence, and DNA sequencing, resulting in the strains H06537 (*ura3(R235A)*) and H06700 (*URA3*).

The appropriate Arg, His, or Thr mutations were introduced into plasmids pPact1-lacZ, YCpU-Pgal-ARG3, pGALOTC, YCpU-Pgal-ASN1, pMAM71, via site-directed mutagenesis performed by Keyclone Technologies (San Marcos, CA). To introduce the R249S mutant of p53 into a yeast plasmid, pLT11, a low-copy *HIS3*-marked plasmid encoding p53(V272M) under control of the yeast *ADH1* promoter (150), was digested with *Sna*BI and *Eco*RI to excise the region including codons 249 and 272, and co-transformed into yeast cells with uncut plasmid pCMV-Neo-Bam p53(R249S), a gift from Bert Vogelstein (Addgene plasmid #16438; http://n2t.net/addgene:16438; RRID:Addgene_16438) (151). WT p53 in the same vector backbone was created by digestion with *Sna*BI and *Stu*I of the low-copy *HIS3*-marked plasmid pTW456 (150) and co-transformation into yeast cells with pMAM38b, a *LEU2*-marked plasmid encoding WT p53. (The WT p53 sequence in pMAM38 was derived from plasmid p663/LR78 (152).) Transformants arising on medium lacking histidine were rescued to *E. coli* and confirmed by sequencing with appropriate primers. The R249S plasmid, called pMAM71, was subjected to site-directed mutagenesis to introduce the H168A or T123A and H168A mutations by Keyclone Technologies, generating plasmids pMAM70 and pMAM113, respectively. Deletion of the mitochondrial import sequence of HsOTC was performed by co-transforming into WT (BY4741) yeast cells *Bam*HI-digested pGALOTC and a double-stranded DNA molecule made by annealing oligonucleotides newhOTCdelNfw (5’ TCCTTTACACAATTAAAAGAAGATGCTGTTTAATCTGAGGGACCTTCTCACTCTAAAAAACTTTACCGGAGAAGAAATTA) and hOTCdelNre (5’TAATTTCTTCTCCGGTAAAGTTTTTTAGAGTGAGAAGGTCCCTCAGATTAAACAGCATCTTCTTTTAATTGTGTAAAGGA). Transformants arising on medium lacking uracil were rescued to *E. coli* and confirmed by PCR and sequencing with appropriate primers.

### Media and additives

*S. cerevisiae* rich growth medium was YPD (1% yeast extract (#Y20020, Research Products International, Mount Prospect, IL), 2% peptone (#P20241, RPI), 2% dextrose (#G32045, RPI)). Supplementation with extra nutrients to test for defects in nutrient uptake was done by adding Leu, His, and uracil to 0.1mg/mL, 0.05mg/mL, and 0.1mg/mL, respectively. Synthetic complete growth medium was based on YC (0.1 g/L Arg, Leu, Lys, Thr, Trp and uracil; 0.05 g/L Asp, His, Ile, Met, Phe, Pro, Ser, Tyr and Val; 0.01 g/L adenine; 1.7 g/L Yeast Nitrogen Base without amino acids or ammonium sulfate; 5 g/L ammonium sulfate; 2% dextrose or galactose) with individual components (from Sigma Aldrich, St. Louis, MO, or RPI) eliminated as appropriate for plasmid selection. For solid media, agar (#A20030, RPI Corp.) was added to 2%. For counterselection against *URA3*, 5-fluoro-orotic acid monohydrate (#F5050, United States Biological, Salem, MA) was added to modified YC medium with 50 µg/mL uracil and 0.1% FOA, for experiments with p53 mutants, or 20 µg/mL uracil and 400 µg/mL FOA for experiments with Ura3 mutants. Nourseothricin (#N5375-74, US Biological) was added to YPD at 100 *μ*g/mL. Hygromycin B (#H-270-1, Gold Biotechnology) was added to YPD at 200 *μ*g/mL. GdnHCl (#G4505, Sigma Aldrich, St. Louis, MO or #G49000, RPI), aminoguanidine HCl (#sc-202931, Santa Cruz Biotechnology, Santa Cruz, CA), N-ethylguanidine HCl (#sc-269833, Santa Cruz), metformin hydrochloride (#M2009, TCI Chemicals), urea (#1610730, Bio-Rad Laboratories, Inc.), TMAO (#AAA1491614, Alfa Aesar), were dissolved in water. MG132 (#10012628, Cayman Chemical Company) was dissolved in ethanol to make a 75 mM stock solution. Medium for MG132 contained, per L: 20 g agar, 20 g dextrose, 0.017 g Yeast Nitrogen Base without amino acids or ammonium sulfate, 1 g Pro, 0.1 g Leu, 0.1 g uracil, 0.05 g His, 0.05 g Met, 0.03 g SDS. Rich medium for *E. coli* was LB (per L: 5 g yeast extract, 10 g tryptone, 10 g sodium chloride), supplemented with carbenicillin or ampicillin and, as appropriate, X-gal (Fisher BioReagents #bp1615-1) from a stock dissolved in dimethylformamide (for experiments with strain RS302) or DMSO (for experiments with DH5*α*).

### Kinetic β-gal assay

We grew 5-mL LB+ampicillin cultures with DH5*α* carrying pACT1-lacZ or its R599A derivative to mid-log phase at 37°C, then added 8 µL of X-gal (20 mg/mL in DMSO) and 5 µL of chloramphenicol (Sigma-Aldrich # C0378, 34 mg/mL dissolved in ethanol). After time intervals of 0, 10, 20, 50, 100, 120 or 150 minutes, a 500-µL aliquot of cells was removed and pelleted, then resuspended in 250 µL of 50 mM Tris-HCl, 10 mM EDTA, pH 8.0, 100 µg/mL RNase A, followed by addition of 250 µL 200 mM sodium hydroxide, 1% SDS to lyse the cells. 100-µL aliquots of this lysate were transferred to the wells of a flat-bottom 96-well microplate. Absorbance at 615 nm was measured with a BioTek Cytation 3 plate reader. Plasmid was isolated from the remainder of the original 5-mL culture of the R599A culture using standard methods and digested with *Pvu*I-HF (New England Biolabs #R3151) according to manufacturer’s instructions. Digested DNA was separated by 1% agarose gel electrophoresis in Tris-acetate EDTA buffer and visualized with ethidium bromide.

### Screening the TS collections

We obtained the Hieter collection as a set of seven plates with colonies on solid medium from the lab of Mark Johnston (University of Colorado Anschutz Medical Campus). A ROTOR bench-top robot (Singer Instruments, Somerset, England) was used to duplicate the colonies to YPD. After incubation at RT, 531 of the 604 mutants formed colonies. These were then pinned by the robot in 384-colony format to two YPD plates and two YPD plates containing 3 mM GdnHCl. One of each type was incubated at RT and the other at 37°C. Growth was scored after 6 days incubation. Unfortunately, photos of these plates were lost, so we do not know the identity of the 73 mutants that did not grow even on YPD at RT. Candidate mutants showing rescue or sensitivity to GdnHCl were streaked to new plates for re-testing.

For GdnHCl rescue, as with the Hieter collection, seven plates of the Boone collection in 96-well format, this time in liquid form, were first robotically transferred to YPD at RT. The resulting colonies were condensed to 384-well format on four YPD plates, and this was done in duplicate for each of three conditions: RT without GdnHCl, 37°C without GdnHCl, and 37°C with 3 mM GdnHCl. After 5 days at the indicated temperature, plates were screened visually to compare colony sizes. We note that a single mutant from the Boone collection that we scored as being rescued by GdnHCl, *erg26-1*, is also represented in the Hieter collection, albeit in a slightly different genetic background. Why it was not identified in the Hieter collection screen is unclear. For urea, DMSO, and TMAO rescue, the colonies arising on YPD after transfer from liquid stocks were pinned in quadruplicate for each of four conditions: YPD at RT and 37°C, and YPD with the selected concentration of experimental small molecule (10 mM urea, 5% DMSO, or 0.5 M TMAO) at RT and 37°C. After 3 days at the indicated temperature, plates were screened visually to compare colony sizes.

### Sequence alignments

Multiple protein sequence alignments were performed using the COBALT tool at the NCBI server (https://www.ncbi.nlm.nih.gov/tools/cobalt/) or, for sequences of yeast species not available via NCBI, using the “Fungal Alignment” function at the Yeast Genome Database (153). In some cases, sequences obtained via the Yeast Genome Database were aligned with other sequences via COBALT.

### DNA sequencing

Genomic DNA from strains *pga1-ts* (H06474), “*cdc20-1*” (the strain found in the well of the Boone TS collection where the *cdc20-1* was supposed to be), *cdc20-2*, *cdc20-3*, *stu1-5* (H06573), the strain with WT “recoded” *URA3* (H06700), or the *ura3(R235A)* strain (H06701) was isolated as described previously and used as template in a Phusion (New England Biolabs, Ipswich, MA) or Taq DNA Polymerase ((New England Biolabs, Ipswich, MA) PCR reaction with primers flanking to or within the coding sequences, according to the polymerase manufacturer’s instructions. These were: 5’PGA1fw (5’ TGTTGACCCTTGGTATTCGC), 3’PGA1re (5’ CCAAAGGGGGCTCAGGCTTC), cdc20_fwd (5’ ATCAGGCCTAGGACTAATCTGCCTTGC), cdc20_rev (5’ CTTTAACATGTGATGAGTTATATCTTGATG), STU1midre (5’GATAACTCATCAGTCAAGTCG), STU1-Nmidfw (5’GAATTTGTTGGCACCGTTCC), STU1-Nmidre (5’CGCTGGGTCTTTACTGCTC), 5’STU1fw (5’GCTGGCAATATAAACACAAGT), 5’URA3fw (5’GGCTGTGGTTTCAGGGTCCAT), 3’URA3re (5’GTCATTATAGAAATCATTACG). Following treatment with alkaline phosphatase (#EF0651) and Exonuclease I (#EN0581, Thermo Scientific Fermentas, Waltham, MA), the purified PCR product was directly sequenced at the sequencing facility of the Barbara Davis Center for Childhood Obesity or Quintara Bioscience, Inc. Additional primers used for sequencing were: midPGA1fw (5’ GTCTTGTCTGACTCGTATCC), midPGA1re (5’ CTGACTCATTGCTTCTCCCG), midURA3re (5’TGCATTCGTAATGTCTGCCC), midURA3fw (5’AAATTGCAGTACTCTGCGGG).

### Fluorescence microscopy

For each sample, a 1-mL aliquot of a 5-mL YPD cultures was pelleted in a microcentrifuge and resuspended in 200 µL of water, then 3 µL was transferred to a 1-mm-thick 1% agarose pad and after the liquid was absorbed, a #1.5 18 mm x 18 mm coverslip was placed on top. Images were captured using an EVOSfl all-in-one microscope (Advanced Microscopy Group, Mill Creek, Washington) using a 60X objective and RFP and GFP LED/filter cubes, as described previously (154). Cytoplasmic fluorescence was quantified using a 6-pixel-diameter circular region, as described previously (155).

## DATA AVAILABILITY

All data are contained within the manuscript.

## SUPPORTING INFORMATION

This article contains supporting information.

## ACKNOWLEDGEMENTS

We thank J.P. Darling-Munson for technical assistance with strain construction, Yoichi Noda (University of Tokyo, Bunkyō-ku) for *pga1* TS strains, Onn Brandman (Stanford University) for the Hsf1 reporter strain, Mark Johnston (University of Colorado Anschutz Medical Campus) for access to the Hieter TS and deletion strain collections and the ROTOR machine, Jay Hesselberth (University of Colorado Anschutz Medical Campus) for access to the Boone TS collection, Bob Sclafani (University of Colorado Anschutz Medical Campus) for the S288C, *erg6*Δ and *mcm3-1* strains, Michael Yeager (University of Colorado Anschutz Medical Campus) for use of the CellASIC temperature monitor, Eric Ross (Colorado State University) for pointing out the Hsp104-independence of [*ISP*^+^] curing by GdnHCl, Joe Nickels (Drexel University College of Medicine) for pointing out the suppression of the TS phenotype of the *erg26-1* mutant, Peter Kaiser (University of California Irvine School of Medicine) for locating and sending the pLT11 and pTW456 plasmids, Stan Fields (University of Washington) for pACT1-lacZ plasmid, and Galit Lahav (Harvard Medical School) for the p53 plasmid p663.

## FUNDING AND ADDITIONAL INFORMATION

This work was supported by the National Institute of General Medical Sciences of the National Institutes of Health under Award R00GM086603 and by a Rare Disease Competition Technology Prize from the BeHEARD Initiative of the Rare Genomics Institute. The content is solely the responsibility of the authors and does not necessarily represent the official views of the National Institutes of Health.

## FOOTNOTES

**Abbreviations:** GdnHCl, guanidine hydrochloride; Gdm, guanidinium; TMAO, Trimethylamine N-oxide; TS, temperature-sensitive; RT, room temperature

---

**Supplementary Figure 1.**
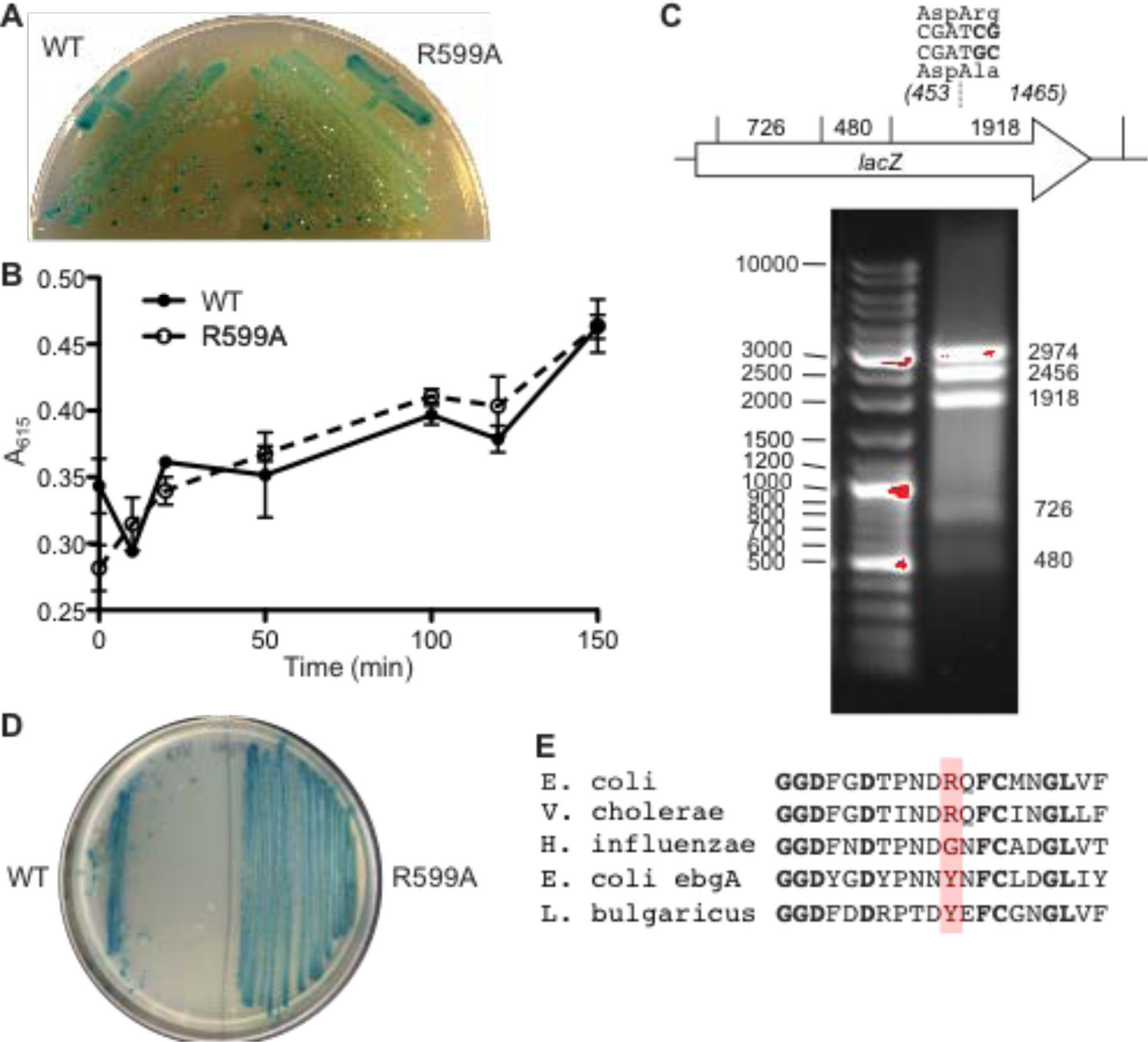
Arg 599 in *E. coli* β-gal is dispensable for activity *in vivo*. (A) *E. coli* cells of strain DH5*α*, carrying the *lacZΔM15* allele at the endogenous locus, were transformed with plasmids carrying WT *lacZ* or the R599A mutant allele under control of the yeast *ACT1* promoter and streaked on medium containing the chromogenic β-gal substrate X-gal at 32 µg/mL. Plates were incubated overnight at 37°C prior to imaging. (B) β-gal activity in cells as in (A) but grown in liquid culture selective for the plasmids, and then lysed at the indicated timepoints following the addition of X-gal and chloramphenicol (to halt new translation). Cleaved X-gal at each timepoint was measured in triplicate by absorbance at 615 nm after the indicated times following X-gal addition. Points show the mean, error bars are standard error of the mean. (C) Plasmid was isolated from the remainder of the R599A culture used in (B) and digested with *Pvu*I before separation by agarose gel electrophoresis. Band sizes are shown in basepairs. Red color indicates saturated pixels. The *Pvu*I recognition site overlapping with the Arg 599 codon is shown with a map (not to scale) of *Pvu*I sites in *lacZ*. DNA ladder was Thermo Scientific GeneRuler DNA Ladder Mix (#SM0331). (D) As in (A) but with the *E. coli* strain RS302, which carries the Δ(*lacIPOZ)C29* allele at the endogenous locus, and with X-gal at 200 µg/mL. (E) Protein sequence alignment of a portion of the indicated β-gal homologs, with *E. coli* Arg 599 highlighted in red and identical residues in bold.

**Supplementary Figure 2.**
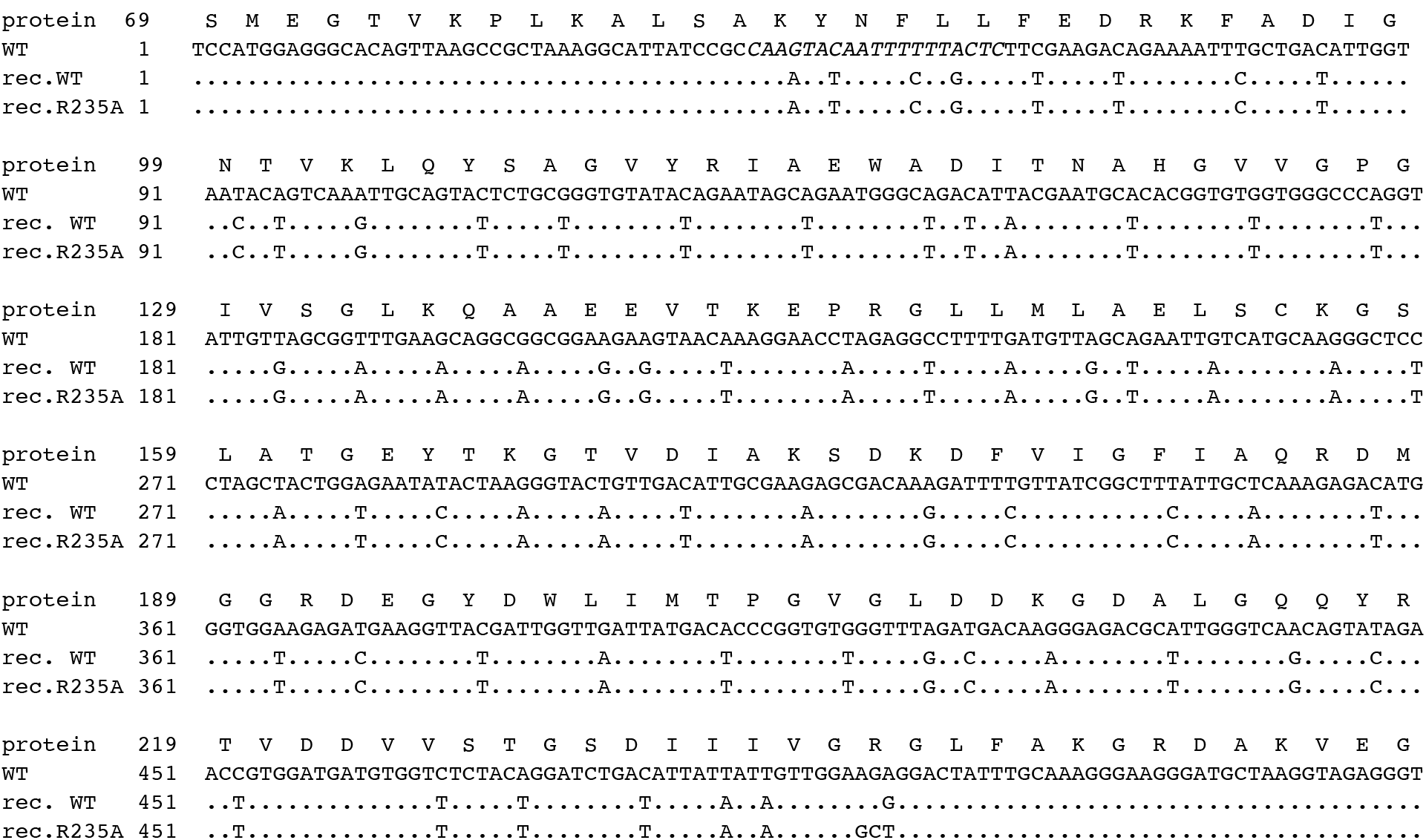
Sequence of “recoded” *URA3* genes used for CRISPR-Cas9-based integration into the yeast genome. “protein”, the amino acid sequence of WT Ura3 starting with residue “WT”, a portion of the DNA sequence of WT *URA3* corresponding to the synthetic sequences below that were used as donor templates for recombination. The sequence corresponding to the guide RNA targeting Cas9 cleavage is italicized. Periods indicate no change in the donor sequences relative to WT.

**Supplementary Figure 3.**
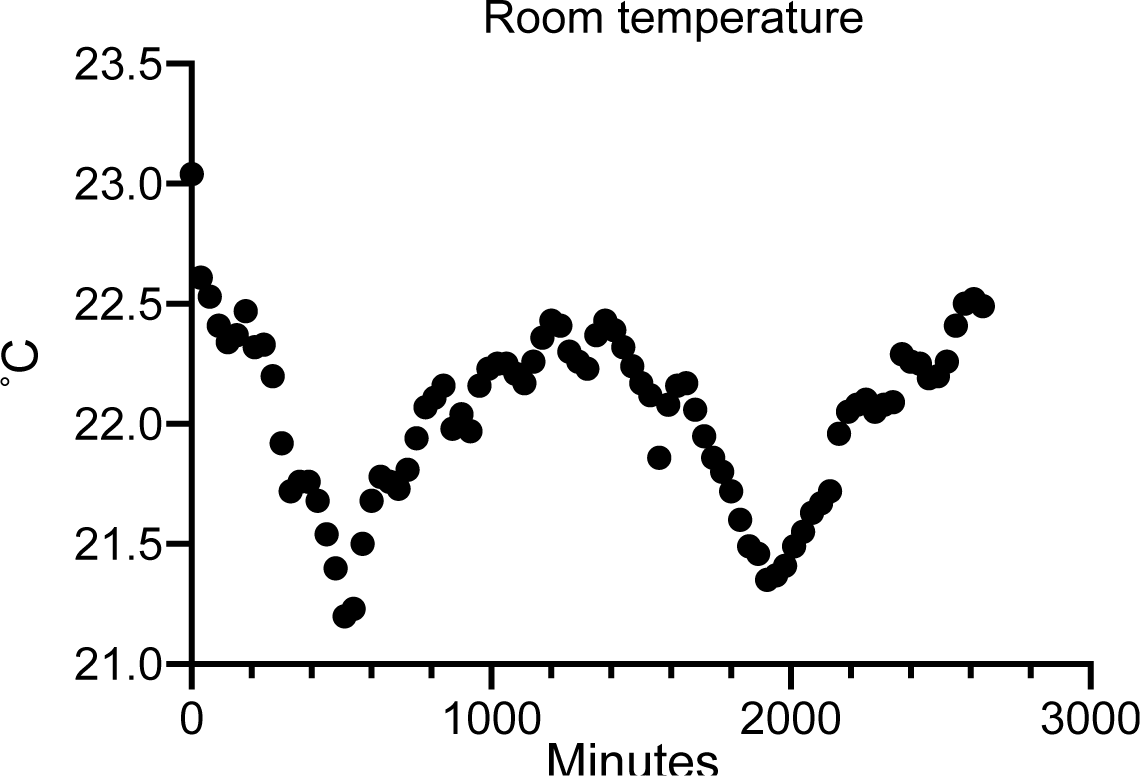
Defining room temperature. Temperature over a nearly 48-hour period was recorded at the location where the majority of the room temperature incubation steps were performed using a CellASIC ONIX™ temperature calibration plate and DirecTemp™ temperature monitoring software.

**Supplementary Figure 4.**
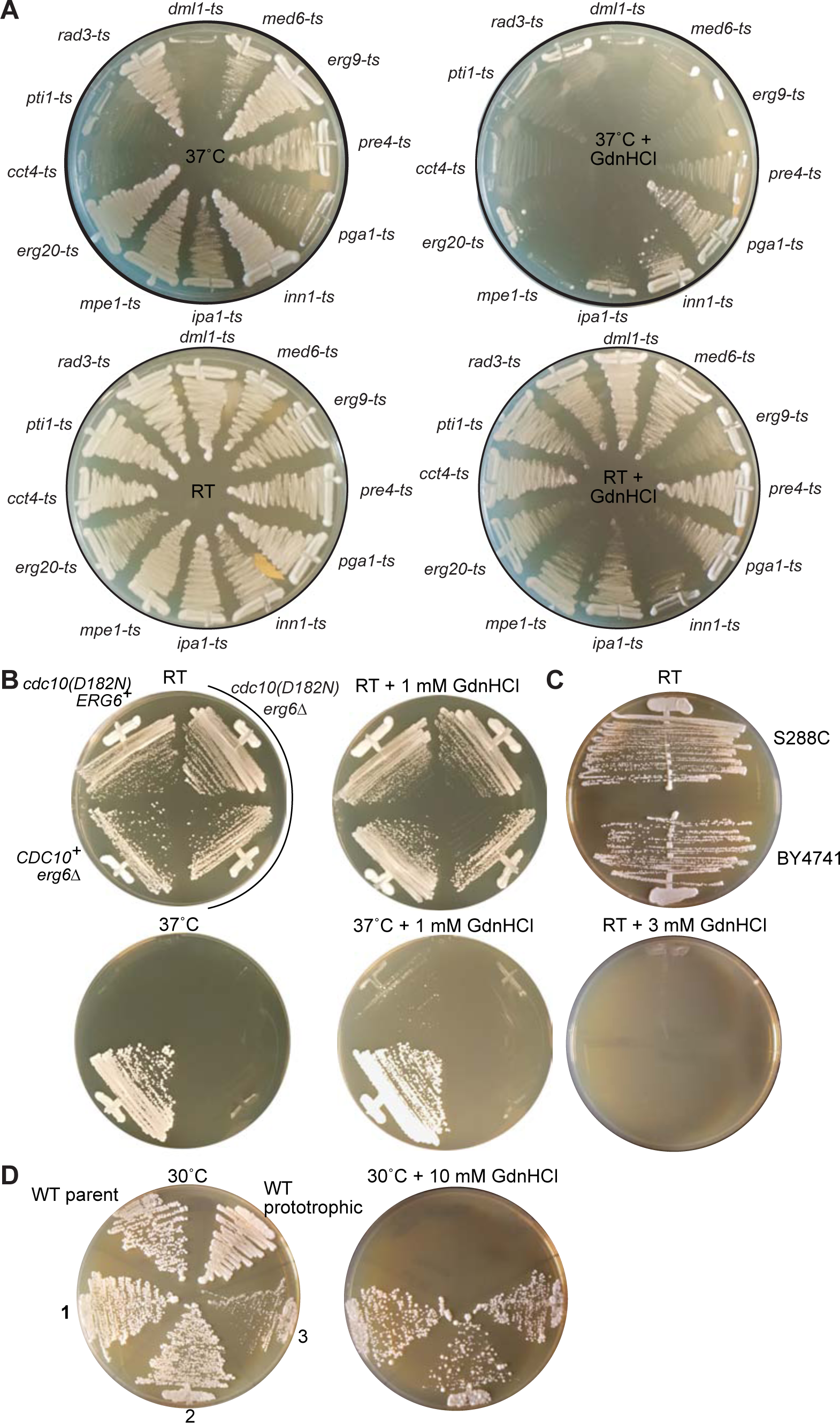
**GdnHCl sensitivity in WT, TS, and *erg6*Δ mutants. (**A) Rich (YPD) plates with or without 3 mM GdnHCl were streaked with TS mutant strains of the indicated genotypes from the Hieter TS collection and incubated at room temperature (“RT”) or 37°C. (B) Rich (YPD) media plates with or without 1 mM GdnHCl were streaked with CBY06147 (“*cdc10(D182N) ERG6^+^*”), H06534 (“*CDC10^+^ erg6*Δ”), and H06537 and another *cdc10(D182N) erg6*Δ haploid clone from the same cross that was not saved as a freezer stock. The plates were then incubated at the indicated temperatures. (C) S288C, a prototrophic WT strain, and BY4741, a strain auxotrophic for histidine, leucine, methionine and uracil but otherwise WT, were streaked to rich (YPD) medium with or without 3 mM GdnHCl and incubated at room temperature (“RT”). (D) Three spontaneously arising GdnHCl-resistant clones (strains H06744, H06745, and H06746, numbered “1”, “2”, and “3”, respectively) were streaked alongside their parent strain, BY4741 (“WT parent”), and S288C (“WT prototrophic”) on rich (YPD) plates with or without 10 mM GdnHCl and incubated at 30°C for 2 or 6 days, respectively, prior to imaging.

**Supplementary Figure 5.**
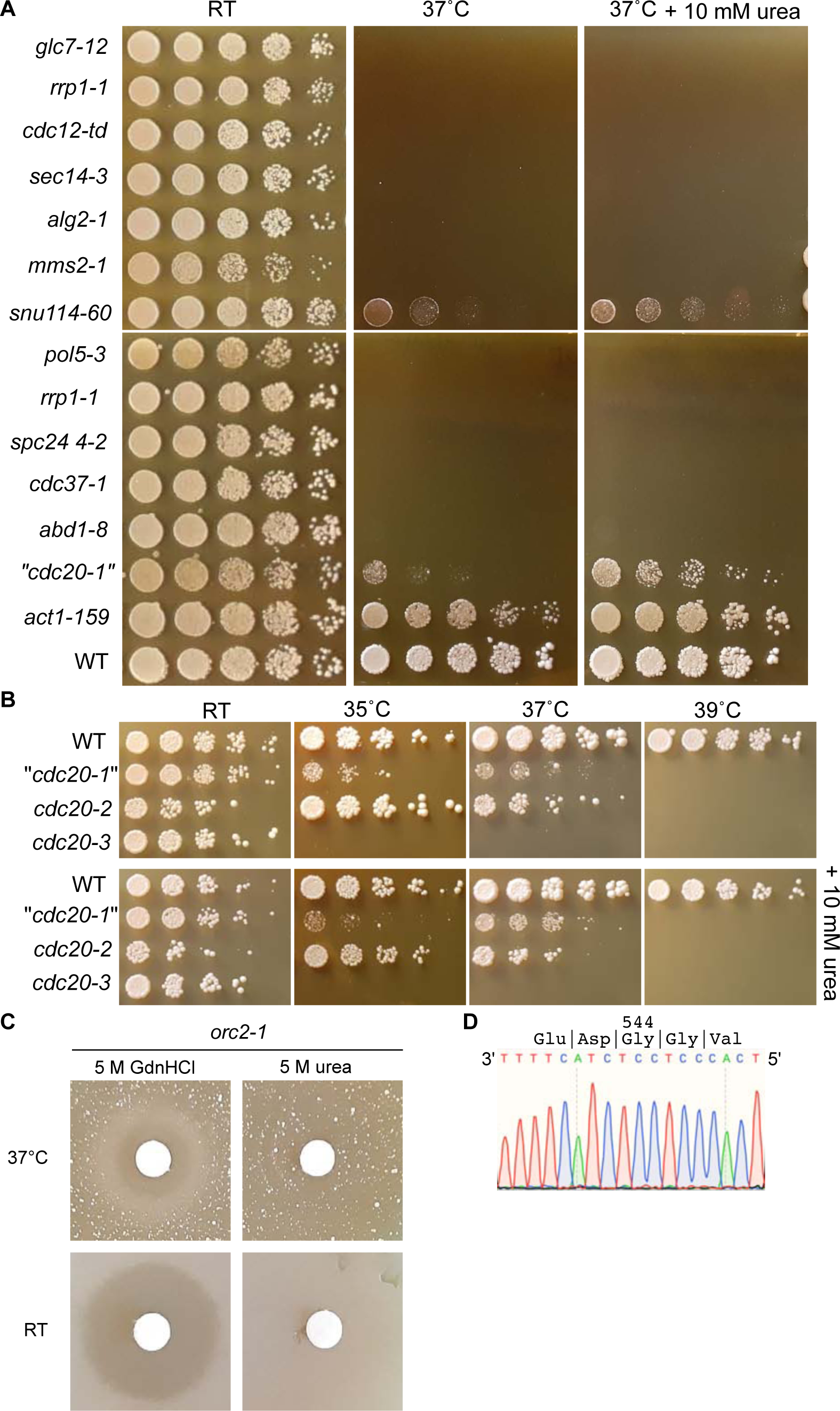
Failure of urea to rescue TS mutants. (A) Mutants from the Boone TS collection scored as possible examples of urea rescue based on robotic screening were serially diluted and spotted on rich (YPD) medium with or without 10 mM urea and incubated at the indicated temperatures. “WT”, BY4741. Note that the bottom row of images is of the same plates as the top, but was cropped and moved to preserve figure space. “*cdc20-1*” is the strain isolated from the well in the TS collection that should have contained the *cdc20-1* mutant, but subsequent PCR and sequencing of the *CDC20* coding region revealed WT sequence at the location of the Gly 544 codon that is mutated in *cdc20-1* (see panel D). (B) As in (A) but with the “*cdc20-1*”, *cdc20-2*, and *cdc20-3* mutants from the Boone TS collection and incubated at additional temperatures. (C) A lawn of *orc2-1* cells (strain CBY08224) was spread on rich (YPD) plates and a paper disc soaked in 5 M GdnHCl or 5 M urea was placed in the center of the plate, after which the plates were incubated at the indicated temperatures. (D) Chromatogram showing dideoxy sequencing results of a PCR product made from genomic DNA of the strain found in the well of the Boone TS collection where the *cdc20-1* mutant (substitution G544R) should have been. Sequencing of PCR products from elsewhere in the *CDC20* coding region of this strain revealed no mutation, whereas sequencing of the same products from the *cdc20-2* and *cdc20-3* strains revealed mutations causing substitutions G360S and E494K, respectively (data not shown).

**Supplementary Figure 6.**
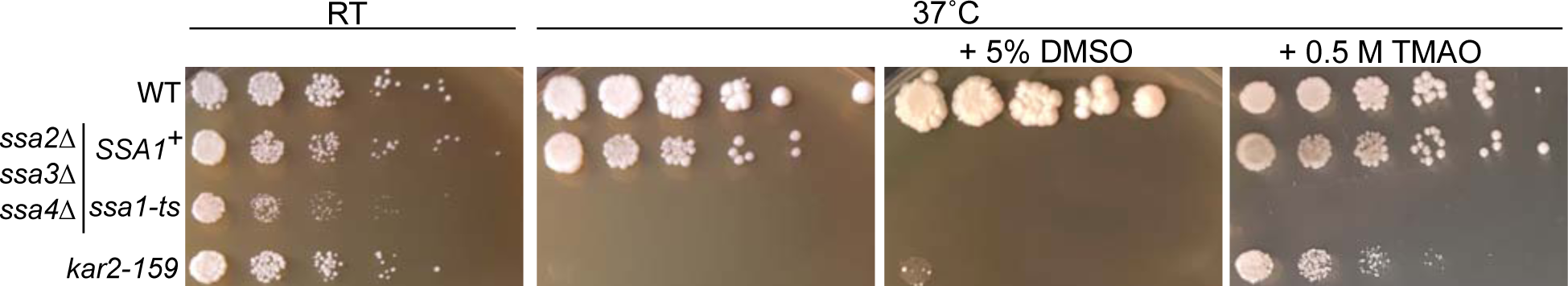
DMSO or TMAO cannot replace cytosolic Hsp70 function. Dilution series spotted on rich (YPD) medium with or without 5% DMSO or 0.5 M TMAO, indicated at the indicated temperatures. The *kar2-159* TS mutant is strain CBY07833 and its WT parent is BY4741. The other two strains, from a distinct strain background, have deletions of three of the four cytosolic SSA-family Hsp70 chaperones (*ssa2Δ ssa3Δ ssa4*Δ) and at the *SSA1* locus carry either a WT allele (“*SSA1^+^*”, strain ECY487) or a TS allele (“*ssa1-ts*”, strain ECY567).

**Supplementary Figure 7.**
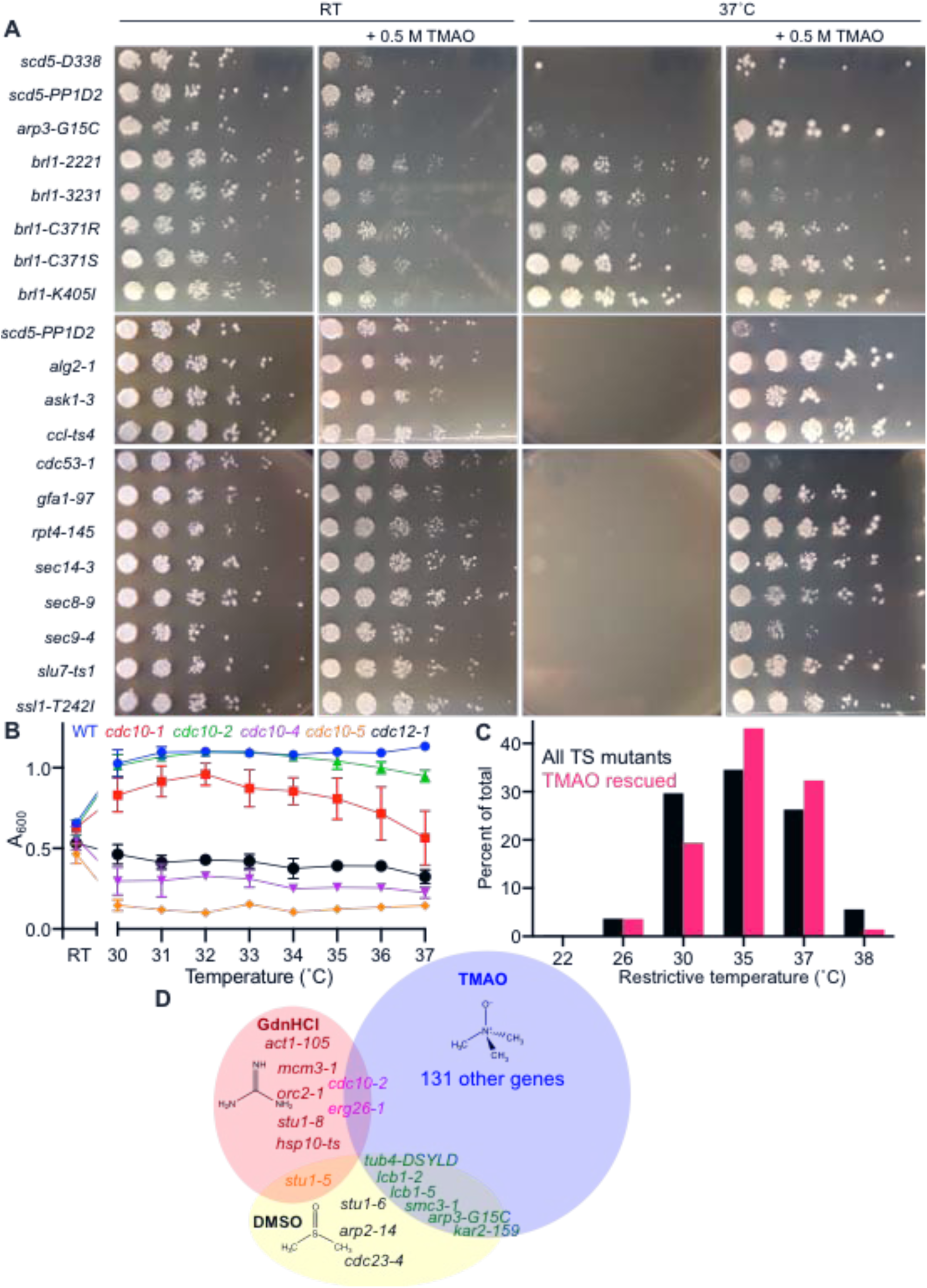
Rescue of TS mutants by TMAO is influenced by restrictive temperature but there is no strict temperature threshold. (A) Strains from the Boone TS collection were serially diluted and spotted on rich (YPD) medium with or without 0.5 M TMAO and incubated at the indicated temperatures. The top row of images represents an independent experiment from the bottom two rows, which were plated at the same time. Note that the *scd5-PP1D2* mutant, identified as rescued by TMAO in the initial robotized screen, showed rescue in only one of the experiments shown here. (B) Cells of the indicated genotypes were suspended in 100 µL rich medium (YPD) in PCR tubes and incubated in a thermal cycler at the indicated temperatures for 20 hours, then the cultures were transferred to a 96-well plate and the optical density was measured at 600 nm. Two technical replicates were performed for each strain; points represent the mean and error bars are standard deviation. Strains were: BY4741 (“WT”), CBY06147 (“*cdc10-1*”), CBY06420 (“*cdc10-2*”), CBY06421 (“*cdc10-4*”), CBY06424 (“*cdc10-5*”), and CBY05110 (“*cdc12-1*”). (C) Histogram of the first temperature listed (140) (see Table S1) as the restrictive temperature for each TS mutant strain from the Boone collection (“All TS mutants”, *n* = 680) and for the mutants from this collection scored as showing some evidence of rescue by TMAO (“TMAO rescued, *n* = 139). (D) Venn diagram of overlap between TS mutants from the Boone collection identified as being rescued by the indicated small molecule. Shape sizes are not to scale.

**Supplementary Figure 8.**
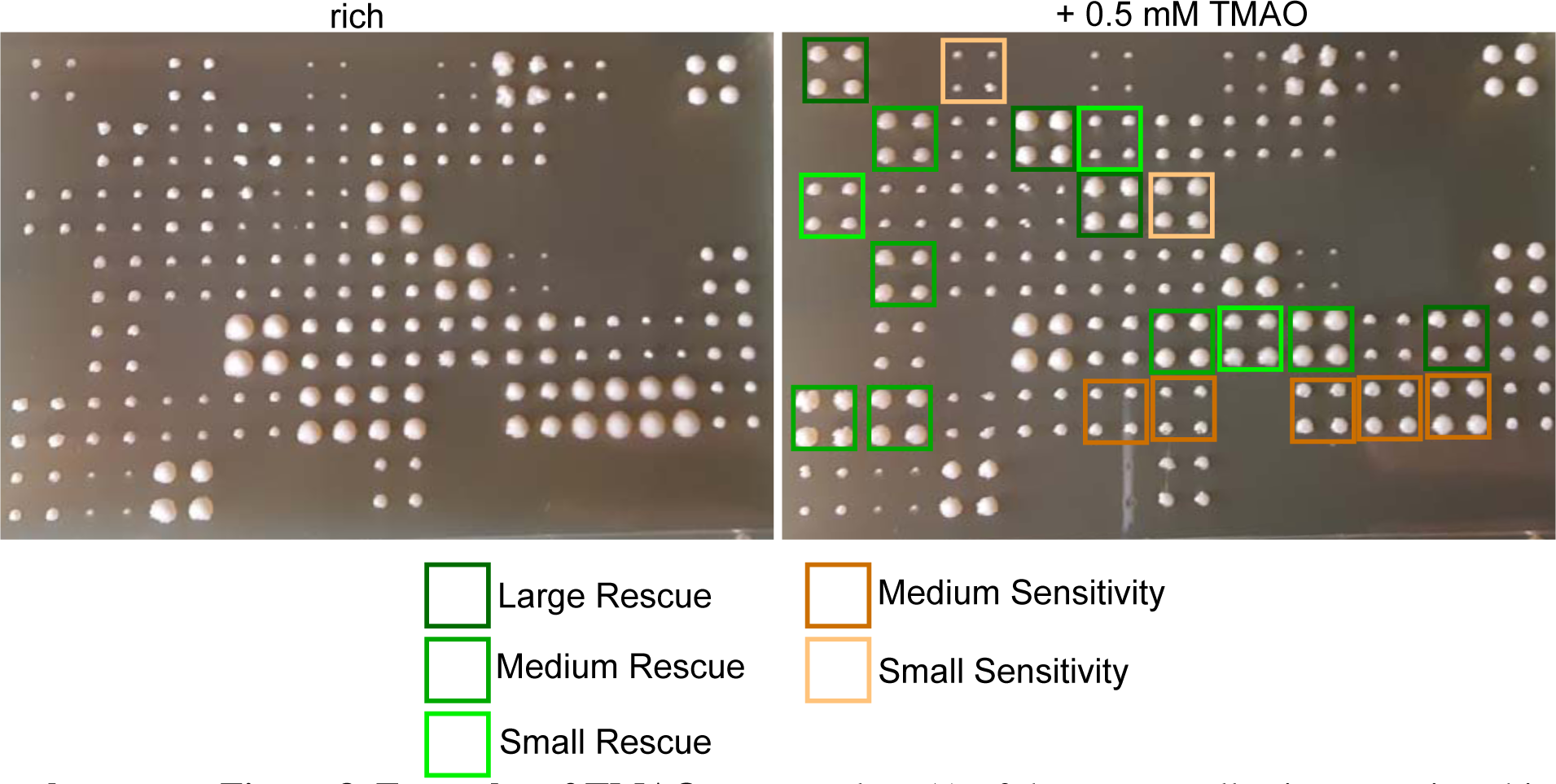
Examples of TMAO rescue. Plate 5 A of the Boone collection was pinned in quadruplicate to rich (YPD) medium with or without 0.5 mM TMAO and the plates were incubated at 37°C prior to imaging.

